# Loss of developmentally derived Irf8+ macrophages promotes hyperinnervation and arrhythmia in the adult zebrafish heart

**DOI:** 10.1101/2024.04.17.589909

**Authors:** Shannon E. Paquette, Cliff I. Oduor, Amy Gaulke, Sabina Stefan, Peter Bronk, Vanny Dafonseca, Nikolai Barulin, Cadence Lee, Rachel Carley, Alan R. Morrison, Bum-Rak Choi, Jeffrey A. Bailey, Jessica S. Plavicki

## Abstract

Recent developments in cardiac macrophage biology have broadened our understanding of the critical functions of macrophages in the heart. As a result, there is further interest in understanding the independent contributions of distinct subsets of macrophage to cardiac development and function. Here, we demonstrate that genetic loss of interferon regulatory factor 8 (Irf8)-positive embryonic-derived macrophages significantly disrupts cardiac conduction, chamber function, and innervation in adult zebrafish. At 4 months post-fertilization (mpf), homozygous *irf8^st96/st96^* mutants have significantly shortened atrial action potential duration and significant differential expression of genes involved in cardiac contraction. Functional *in vivo* assessments via electro- and echocardiograms at 12 mpf reveal that *irf8* mutants are arrhythmogenic and exhibit diastolic dysfunction and ventricular stiffening. To identify the molecular drivers of the functional disturbances in *irf8* null zebrafish, we perform single cell RNA sequencing and immunohistochemistry, which reveal increased leukocyte infiltration, epicardial activation, mesenchymal gene expression, and fibrosis. *Irf8* null hearts are also hyperinnervated and have aberrant axonal patterning, a phenotype not previously assessed in the context of cardiac macrophage loss. Gene ontology analysis supports a novel role for activated epicardial-derived cells (EPDCs) in promoting neurogenesis and neuronal remodeling *in vivo*. Together, these data uncover significant cardiac abnormalities following embryonic macrophage loss and expand our knowledge of critical macrophage functions in heart physiology and governing homeostatic heart health.

## INTRODUCTION

Macrophages have garnered much attention in recent years for their vast array of non-canonical functions that extend beyond their historical characterization as sentinel immuno-modulatory cells. These functions can be broadly categorized into organ or cell-specific developmental regulation, adult tissue homeostasis and maintenance, and regeneration^1–10^. Likewise, several reports have identified important novel functions for cardiac macrophages in heart development, homeostasis, electrical modulation, and disease^11–22^. The first wave of cardiac macrophages in mice is detectable around embryonic day (E) 10.5 with further expansion occurring through E14.5^19,23,24^. Lineage tracing experiments have also revealed heterogeneous origins, with >90% of cardiac-seeded macrophages at E16.5 deriving from the yolk-sac, fetal liver, and hemogenic endothelium through endothelial-to-hematopoietic transition (EHT) during primitive and transient definitive hematopoiesis, and the remaining populations derived from the aorta-gonad-mesonephros (AGM) during definitive hematopoiesis^19,23–25^. These seeding events in early development are largely responsible for the local, self-maintaining population seen in the adult heart^23,26^, though aging or loss of this population leads to an increase in recruited monocyte-derived macrophages^23,27^. It is currently unclear the extent to which monocyte-derived macrophages can adopt embryonic-derived macrophage functions.

Evidence indicates that embryonic-derived tissue-resident macrophages are integral to coronary arterial remodeling, maturation, and cardiac tissue repair^11,12,18,24,28^, and that monocyte-derived macrophages may not be able to fully recapitulate these functions. For example, age-dependent susceptibility studies using a diphtheria toxin-induced cardiac injury model revealed that neonatal mouse hearts have a distinct macrophage profile following injury that consists of an expansion of the resident macrophage populations rather than a recruitment of monocytes, as seen in the adult heart post-injury^18^. The fetal hearts had significantly reduced inflammation and scarring compared to adult hearts^18,29,30^, suggesting that resident and recruited macrophages may carry out distinct functions that warrant further investigation. Indeed, there is much interest in understanding the independent contributions of each macrophage population to cardiac development and function.

Macrophages have multiple important functions in adult heart conduction and homeostasis that fall outside of their traditional roles as immune cells. Macrophages were recently discovered to be capable of electrically modulating cardiac activity by coupling to cardiomyocytes at the adult mouse atrioventricular node (AVN) via the gap junction connexin43 (Cx43)^16^. Consistent with these experimental findings, macrophages were found in the adult human AVN, suggesting that they may modulate conduction in the human heart^16^. Macrophages have also been shown to maintain Cx43 phosphorylation and localization within the mouse myocardium through expression of macrophage-derived amphiregulin (AREG)^21^. *Areg*^-/-^ mice had increased incidence of arrhythmias and significantly increased risk of sudden death following right ventricular overload via pulmonary artery banding^21^. Lastly, cardiac macrophages were demonstrated to support mitochondrial homeostasis and energy utilization in the heart by phagocytosing packaged debris, i.e., membrane-surrounded vesicles or exophers, from healthy cardiomyocytes^22^. This important clearance function was necessary to prevent inflammasome activation, autophagic block, and metabolic dysfunction by altered availability of ATP^22^. While these studies expand our appreciation of macrophage functions in the adult heart, they still do not provide clarity as to whether embryonic-derived macrophages are specifically important for adult heart physiology and tissue homeostasis.

In this study, we sought to determine how loss of embryonic-derived macrophages impacted adult heart function and the molecular and cellular landscape using the zebrafish model. The conservation of cardiac regulatory pathways, comparable electrophysiology, and ability to recapitulate human pathophysiology all make zebrafish an attractive model for studying congenital and acquired heart defects^31–33^. Conserved innate and adaptive immune profiles also contribute to the zebrafish’s applicability for immunological studies, including macrophage biology^34–36^. In zebrafish development, macrophage commitment during primitive myelopoiesis was shown to be governed by expression of interferon regulatory factor 8 (Irf8)^37^. Zebrafish with a null mutation in *irf8* lack macrophages for the first ∼7 days of development^38^. This developmental timeline parallels the onset of hematopoietic stem cell (HSC) differentiation within the kidney marrow, the bone marrow equivalent in zebrafish^39^. Therefore, *irf8* mutant zebrafish lack primitive and transient definitive macrophage populations and any *de novo* populations of macrophages are likely derived from blood monocytes.

Using wild type (WT) and homozygous *irf8^st^*^96^*^/st^*^96^ mutant zebrafish, we performed an array of electrical, physiological, transcriptomic, and immunohistological assessments to determine how loss of embryonic-derived macrophages affects the adult heart. Optical mapping and electrocardiogram (ECG) studies reveal that *irf8* mutants have significant electrical abnormalities, which are further supported by echocardiograms revealing diastolic dysfunction and pumping deficiencies. Hearts of *irf8* null zebrafish are also hyperinnervated and have aberrant neuronal patterning. Transcriptomic and immunostaining analyses reveal a novel role for activated epicardial-derived cells (EPDCs) of *irf8* null hearts in neurogenesis, suggesting that stress-responsive EPDCs are involved in neuronal remodeling and the observed arrhythmias. Together, this study defines the importance of embryonic-derived macrophages in cardiac function and in maintaining adult heart homeostasis.

## RESULTS

### Embryonic macrophages modulate cardiac function

To assess whether macrophages carry out non-canonical functions in the developing zebrafish heart, we first investigated whether macrophages had the capacity to influence cardiac activity. While it was previously shown that macrophages are abundant at the AVN in adult murine and human hearts^16^, it is unclear whether macrophages reside at the SAN, the electrical pacemaker that generates impulses to initiate heart contraction. Using the transgenic zebrafish line *Tg*(*mpeg1:EGFP)*^40^ to mark macrophages, we found that macrophages reside at the SA region at the base of the atrium in close association to Isl1^+^ pacemaker cardiomyocytes as early as 4 days post-fertilization (dpf) (Figure S1A-B’). Macrophage presence at the zebrafish SAN was maintained into adulthood (Figure S1C-D’). Using tamoxifen-inducible Csf1r-tdTomato mice, we confirmed that macrophages were present at the SAN in mammalian hearts as early at E16.5 (Figure S1E-E’) and continue residency at the SAN in post-natal hearts (P24; Figure S1F-F’). To address whether macrophages had the potential to influence pacemaker electrical activity, we performed a stain against connexin45 (Cx45), a gap junction protein abundant at the SAN that electrically couples neighboring cells to allow for ion exchange and impulse propagation. Cx45 has been implicated in myocyte-myocyte coupling as well as myocyte-fibroblast coupling^41^, though it is not known whether cardiac macrophages express Cx45. We found that macrophages within the murine SAN do, in fact, express Cx45 with abutting pacemaker cardiomyocytes (Figure S1G-H), indicating that they could play a direct role in nodal electrical balance or function.

Based on the timing of macrophage seeding at the zebrafish and murine SAN, cardiac macrophages in 4 dpf zebrafish and E16.5 fetal mice are presumed to be derived during primitive or transient definitive hematopoiesis^19,23,24,38^, hereafter referred to as embryonic-derived macrophages. In zebrafish, embryonic-derived macrophages can arise from the zebrafish intermediate cell mass (ICM, murine yolk sac equivalent), AGM, or caudal hematopoietic tissue (CHT, murine fetal liver equivalent)^42^ in the first week of development. Considering that embryonic-derived macrophages comprise tissue resident, self-maintaining cardiac macrophage populations^19,23,26^, we asked whether embryonic macrophages could modulate the larval zebrafish heart *in vivo*. To do this, we employed optogenetics to manipulate macrophage membrane potential at 7 dpf and observed the concomitant effects on heart function. Using the Gal4/UAS system to achieve targeted opsin expression, we generated zebrafish with macrophage-specific expression of the light-gated cation channel channelrhodopsin (ChR2)^43^ (*Tg*(*mpeg1:Gal4FF:UAS:ChR2-mCherry*)) (Figure 1A). Heart rate was captured for: (i) 30 seconds prior to stimulation, (ii) 30 seconds during macrophage stimulation with 405 nm light, and (iii) 10 minutes post-stimulation. Stimulation of ChR2+ macrophages resulted in a significant increase in heart rate over ChR2-sibling controls (Figure 1B). Heartbeat also became irregular in the ChR2+ larvae, which was not attributed to blue light stimulation (Supplemental Video 1 versus 2). Because our system for opsin stimulation illuminates a broad area, there was the potential that non-cardiac macrophages were also stimulated. We therefore confirmed stimulated macrophages were not influencing neuronal activity, thereby indirectly modulating heart rate. Indeed, pan-neuronal stimulation of ChR2 (*Tg*(*elavl3:Gal4-VP16;UAS:ChR2-mCherry*) did not result in an increase in heart rate (Figure 1C). Of note, we observed differences in baseline heart rate between the transgenic backgrounds (“No Stim.,” Figure 1B versus 1C). Such phenotypic variations have been noted in other contexts as well and are linked to the high number of transposable elements present in both teleost and mammalian genomes^44^. We hypothesize that this may also be why the ChR2-negative control fish expressing *Tg*(*elavl3:Gal4-VP16)* are responding to the blue light (Figure 1C). Nonetheless, the specificity of increasing heart rate in macrophage-driven over neuron-driven ChR2+ fish strongly supports a role of embryonic macrophages in electrical modulation of the SAN in early in development.

**Figure 1.**
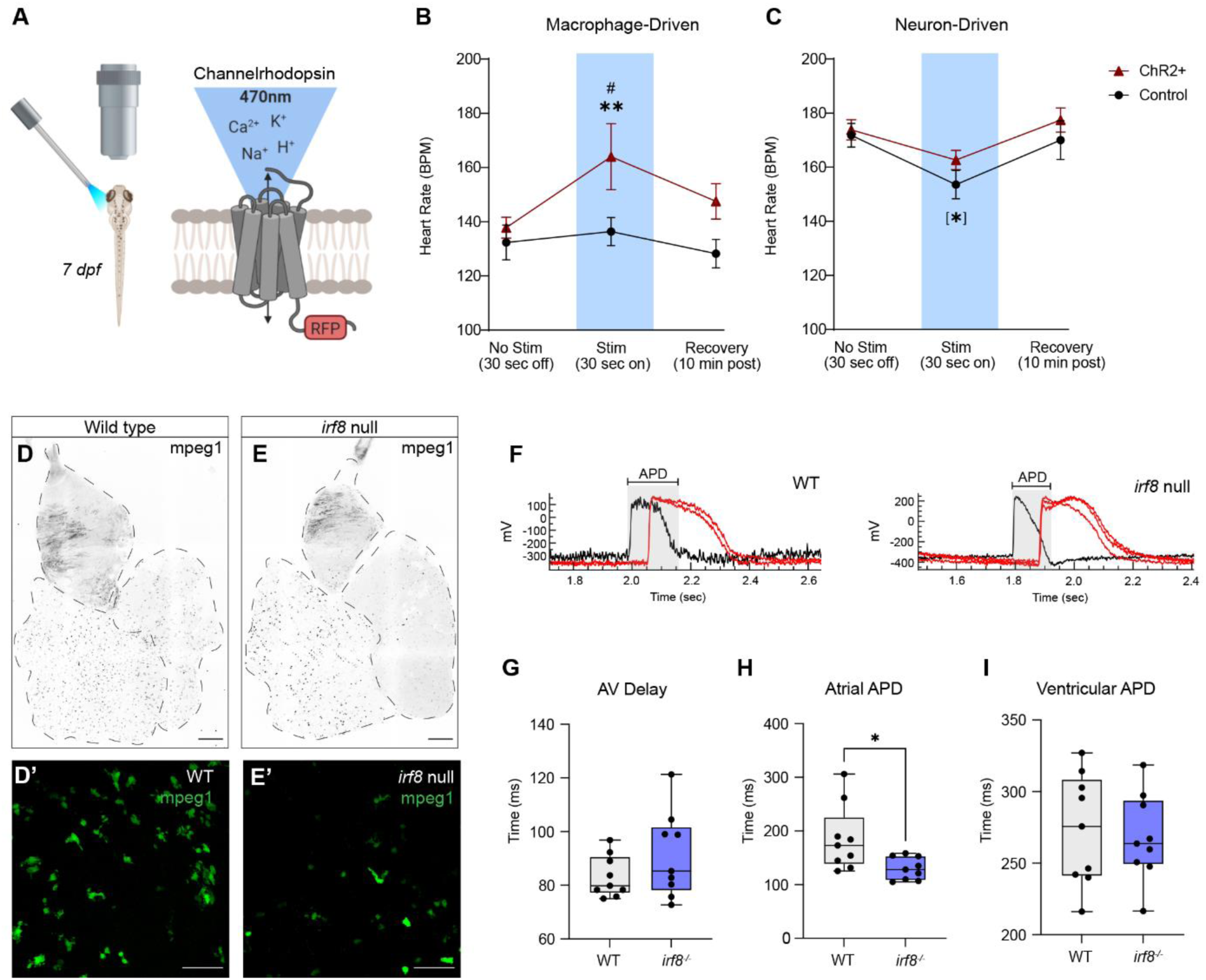
Loss of embryonic macrophages produces electrical and transcriptional signatures of cardiac dysfunction. (A) Cartoon depiction of optogenetic stimulation of channelrhodopsin (ChR2)-positive macrophages with blue light. (B-C) Heart rate of 7 dpf control larvae and larvae with ChR2+ macrophages (B) or neurons (C). Heart rate was determined at three different timepoints using the following stimulation protocol: 30 seconds before stimulation (“No stim.”), 30 seconds during stimulation with 405 nm light (“Stim.”), and 10 minutes post-stimulation (“Recovery”). #p < 0.05, “No Stim.” ChR2+ versus “Stim.” ChR2+; **p < 0.01, “Stim.” Control versus “Stim.” ChR2+. Welch’s t-test. [*]p < 0.05, “No Stim.” Control versus “Stim.” Control. (D-E’) Confocal micrographs of 12 mpf wild type (D-D’) and *irf8* mutant (E-E’) hearts with transgenic expression of GFP-labeled macrophages (mpeg1+). Scale = 200 um (D,E) and 50 um (D’,E’). Images at 10x (D,E) and 40x (D’,E’). (F) Representative optical voltage mapping traces of isolated 4 mpf *irf8* wild type and mutant (null) hearts bathed in 10 µm di-4-ANNEPS. Atrial activation (black); ventricular activation (red). Gray box indicates atrial action potential duration (APD). (G-I) Quantification of AV delay (G), atrial action potential duration (APD) (H), and ventricular APD (I). n = 9 per genotype. Welch’s t-test.

We next sought to determine whether loss of embryonic-derived macrophages impacted adult heart function using *irf8^st^*^96^*^/st^*^96^ mutants^38^. Adult *irf8* mutants had Mpeg1+ cells within the atrium, ventricle, and outflow tract (Figure 1D-E’). Since *mpeg1* is also expressed by B cells in zebrafish^45^, we further confirmed macrophage expression in the heart via *in situ* by probing for both the macrophage-specific gene *mfap4* and B cell-specific gene *pax5*. Indeed, adult *irf8* mutants expressed *mfap4+*/*pax5-* cells in the atrium and ventricle, confirming the presence of macrophages in the absence of Irf8 function (Figure S2). To assess whether cardiac conduction was disrupted in *irf8* mutant fish, we performed optical voltage mapping on isolated 4 months post-fertilization (mpf) zebrafish hearts stained with 10 µM di-4-ANEPPS. Voltage mapping experiments revealed that *irf8* mutants had significantly reduced atrial action potential duration (Figure 1F,H). Whole tissue analysis from 4 mpf single cell RNA (scRNA) sequencing data also revealed multiple significantly differentially expressed pathways involved in heart contraction, circulation, nucleic acid processes, and sarcomere organization (Figure S3). Together, these data suggest that loss of Irf8+ macrophages result in both functional and transcriptional alterations in the adult zebrafish heart.

### *irf8* mutants have increased incidence of arrhythmia post-exercise

We next sought to determine whether adult *irf8* null zebrafish exhibited arrhythmias at baseline and after exercise by performing electrocardiograms (ECGs). At 12 mpf, *irf8* mutants had a significant increase in basal heart rate (+24.5% increase), but there were no other notable differences in the trace measurements (Table S1). To determine if *irf8* mutants were susceptible to developing arrhythmias, we challenged the fish with a swim tunnel exercise assay and immediately followed the physiological challenge with an ECG (Figure 2A-B). Swim tunnels are enclosed chambers with continuous laminar water flow at controllable flow rates that provide important information on bioenergetics, biomechanics, and respirometry^46,47^. For the swim assay, we designed a custom, rack-compatible swim tunnel that achieved laminar flow, had adjustable flow rate, and was equipped with a digital flow meter for monitoring (See ‘Methods’). This innovative swim tunnel design permitted direct use of fresh oxygenated and temperature-controlled facility water from our facility racks. Following a 1-hour light-to-moderate swim challenge, both male and female *irf8* mutants had a significant increase in various arrhythmic events, including premature atrial contractions, spontaneous peaks, sinus arrhythmia, and irregular R wave amplitude (Figure 2C-D; Figure S4). We also observed flutter-like electrical disturbances in the traces that occurred above the baseline noise (Figure 2C). Because fish were anesthetized during the recording, these were considered bona fide electrical events rather than movement artifacts. Interestingly, there were no significant differences in the average P duration, PR interval, RR interval, or QRS interval duration (Table S2), indicating that the electrical anomalies of individual peaks were masked when averaged together. While 0% of wild type females and 28% of wild type males exhibited an arrhythmic event post-swim challenge, arrhythmic events were present in 63% of mutant females and 71% of mutant males and (p < 0.0001; Figure 2D-E). Upon categorizing the major arrhythmic events exhibited by all samples, there is a significant increase in incidence or co-incidence of these arrhythmic events in *irf8* mutants (p *<* 0.0001).

**Figure 2.**
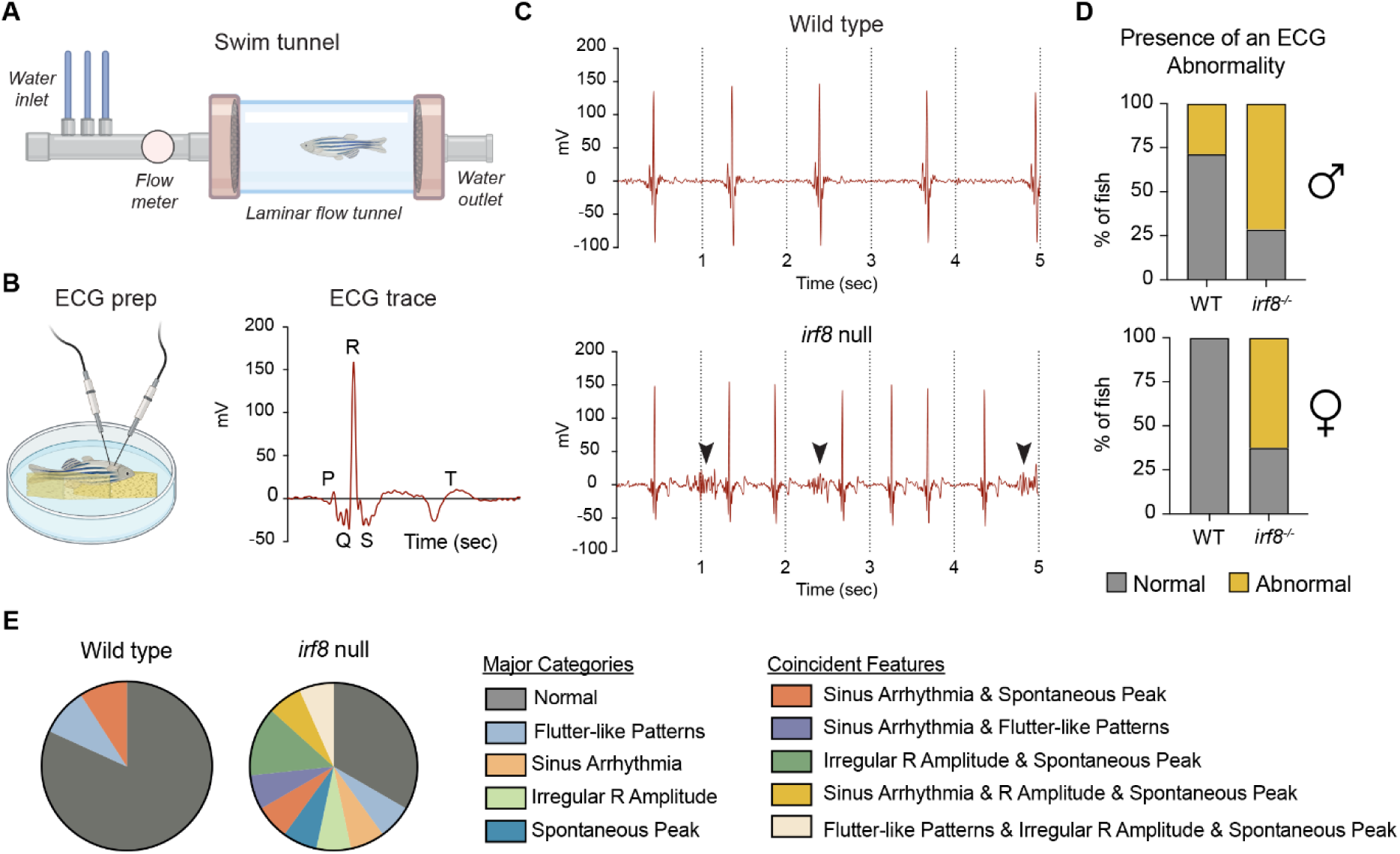
Functional swim tunnel challenge reveals *irf8* mutants are susceptible to arrhythmia. (A) Cartoon depiction of the rack-compatible swim tunnel with metered flow. Following a 5-minute acclimation period within the tunnel, 12 mpf zebrafish were subjected to increasing flow rate intervals for a total of 1 hour. (B) Cartoon depiction of an ECG recording and a representative ECG trace from an anesthetized fish placed ventral-side up in a sponge surrounded by anesthesia solution. (C) Representative ECG traces for wild type and *irf8* mutant zebrafish following the swim assay. Black arrows indicate unique electrical disturbances occurring above any baseline noise. (D) Percentage of fish with an abnormal electrical event during the ECG recording. Male n = 7 per genotype, ****p < 0.0001. Female n = 4-8 per genotype, ****p < 0.0001. Fisher’s exact test. (E) Categorization of ECG abnormalities observed throughout all wild type and *irf8* mutant fish.

### *irf8* mutant hearts have a pumping deficiency as well as age- and sex-specific effects on chamber function

While ECGs provide electrical assessments of the heart, echocardiograms provide important insight into cardiac structure, hemodynamics, and chamber function. We therefore performed high-frequency echocardiography at both 6 mpf and 12 mpf in anesthetized fish (Figure 3A-D). While echocardiograms have been used for many decades both clinically and in mammalian research, only recently has this modality been adapted and standardized for use in adult zebrafish^48,49^. Ultrasound images were acquired in B-mode for ventricular volume, area, fractional shortening, ejection fraction, and stroke volume, as well as in pulsed-wave Doppler to assess AV inflow and ventriculobulbar (VB) outflow. The echocardiographic measurements revealed several age- and sex-specific functional differences in *irf8* null hearts (Table S3). For example, aortic acceleration time (AAT), the time between VB valve opening to peak aortic jet velocity, was significantly prolonged in both 6 mpf *irf8* mutant males and females compared to wild type (+25.0%, p<0.0001 and +33.1%, p<0.0001, respectively), but was not prolonged at 12 mpf (Table S3). Interestingly, *irf8* null AAT durations at 6 mpf were similar to wild type durations at 12 mpf, indicating that (1) *irf8* mutant hearts may exhibit an early aging phenotype and (2) chamber dysfunction occurring earlier in life may pseudonormalize with age, which is also observed in humans^50–52^. There were, however, measurements consistently altered in both sexes and at both timepoints in the *irf8* mutants, including significantly prolonged isovolumic relaxation time (IVRT; Figure 3E), significantly prolonged aortic ejection time (AET; Figure 3F), and significantly reduced pressure within the VB valve (VB Mean Gradient; Figure 3G). Additionally, at 12 mpf, *irf8* mutant males had a moderate reduction (-14.7%; p = 0.195) and females had a significant reduction (-33.7%; p = 0.0073) in fractional shortening (Table S3). Prolonged IVRT, which is the time between VB closure and AV valve opening, is an indicator of diastolic dysfunction and poor myocardial relaxation. Likewise, prolonged AET is associated with diastolic dysfunction, arterial stiffening, aortic stenosis, and all-cause mortality^53^. Low gradient, i.e. pressure, and reduced transaortic velocity within the VB valve can be an indicator of aortic stenosis, though it can also be secondary to other ventricular dysfunctions such as myocardial infarct and fibrosis^54^. Together, echocardiographic measurements therefore revealed that *irf8* mutants have signatures of diastolic dysfunction and ventricular stiffening.

**Figure 3.**
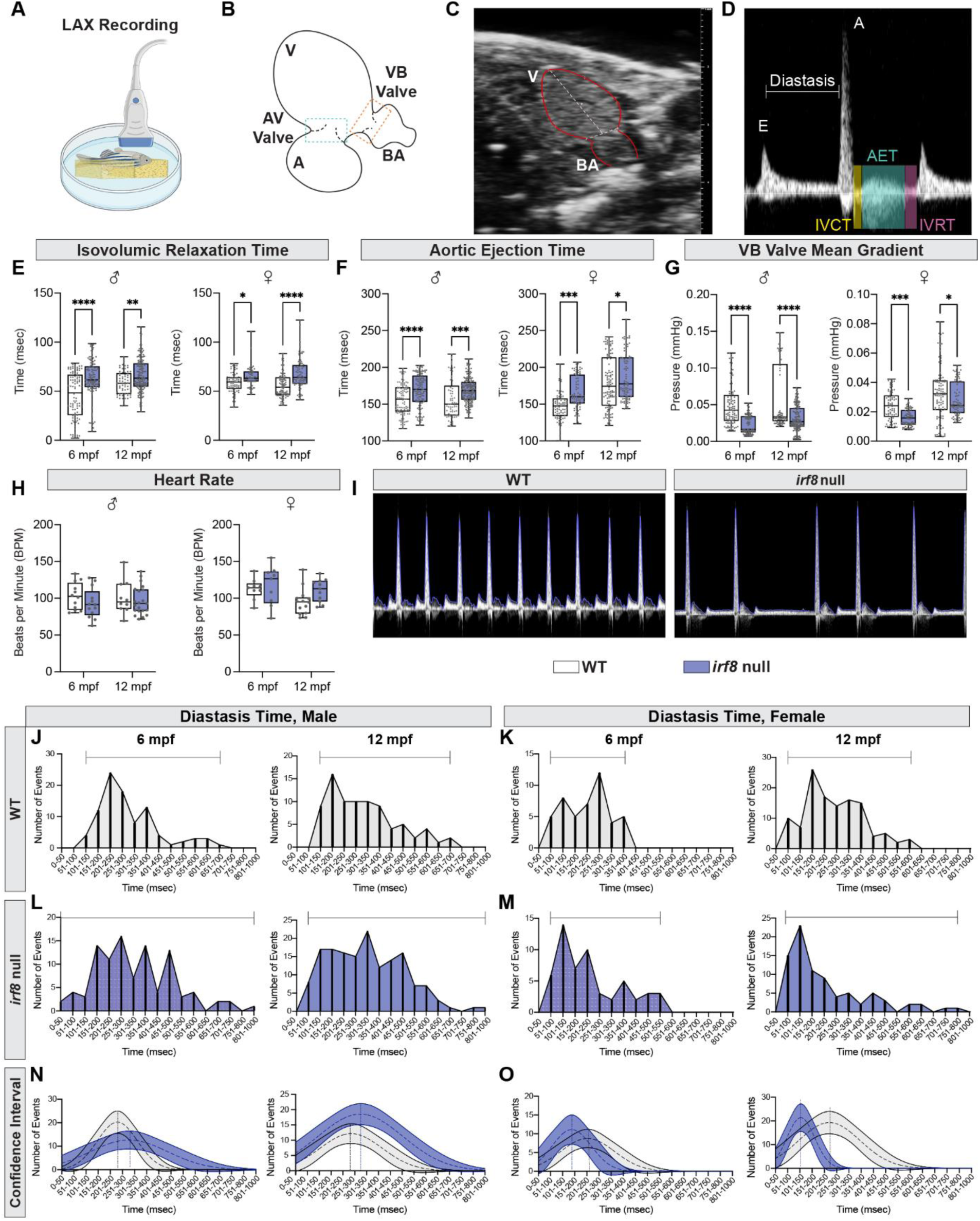
Echocardiographic measurements further support arrhythmicity in *irf8* mutants and identify a pumping deficiency. (A) Cartoon depiction of an adult zebrafish undergoing an echocardiogram with the transducer in the long-axis (LAX) position. Echocardiograms were performed on unchallenged fish. (B) Cartoon of an adult zebrafish heart from the perspective of the echocardiography recording, denoting the atrium (A), atrioventricular (AV) valve, ventriculobulbar (VB) valve, and bulbus arteriosus (BA). The boxes around the AV valve and VB valve depict the gating strategies to determine inward flow and outward flow of blood from the ventricle. (C) Still image from an adult zebrafish echocardiography recording with the ventricle and bulbus arteriosus outlined in red. (D) Representative echocardiogram trace gating on the AV valve with the early filling (E) wave, late filling (A) wave, diastasis time, isovolumic contraction time (IVCT), aortic ejection time (AET), and isovolumic relaxation time (IVRT) labeled. (E-G) Quantifications of IVRT (E), AET (F), and pressure within the VB valve (G) in male and female wild type and *irf8* mutant fish at 6 and 12 mpf. Male n = 10-21 per genotype and timepoint. Female n = 9-14 per genotype and timepoint. Each point represents individual beats per fish. 8-10 beats were analyzed per fish. *p < 0.05; **p < 0.01; ***p < 0.001; ****p < 0.0001. (H) Quantification of heart in male and female wild type and *irf8* mutants. (I) Representative echocardiogram traces from a wild type and *irf8* mutant recording. (J-M) Quantification of diastasis time in wild type males (J) and females (K) at 6 mpf and 12 mpf, as well as diastasis time in *irf8* mutant males (L) and females (M) at 6 mpf and 12 mpf. A Gaussian distribution was created by analyzing 8 beats per sample and grouping diastasis duration in defined increments. Male n = 10-21 per genotype and timepoint. Female n = 9-14 per genotype and timepoint. Horizontal bars represent the range of diastasis duration. (N-O) 95% confidence interval of the average diastasis time durations in wild type (gray) and *irf8* mutant (blue) males (N) and females (O).

According to the echocardiographic analysis, *irf8* mutants did not have a significant difference in heart rate at 6 mpf or 12 mpf (Figure 3H); however, they did exhibit arrhythmia in the form of irregularity in spacing between A waves (Figure 3I), reminiscent of the sinus arrhythmia exhibited in the ECG recordings. It was determined that this observed variability was driven by the diastasis time, which is the time between early filling (E wave) and late filling (A wave) of the ventricle (Figure 3D). At both 6 mpf and 12 mpf, male and female *irf8* mutants demonstrated an increased range of diastasis times within the population (Figure 3J,K versus 3L,M), as well as a shift in the average diastasis time duration compared to wild type fish (Figure 3 N-O). While diastasis time occurred on average from 200 to 300 msec in wild type fish, diastasis time in the mutants ranged from <100 msec to >800 msec in both males (Figure 3L) and females (Figure 3M). Additionally, *irf8* mutant males had prolonged diastasis time between 300 and 350 msec (Figure 3N), while mutant females had shortened diastasis time between 150 and 200 msec (Figure 3O). Unlike humans, where the E/A ratio is ≥ 1, ventricular filling in zebrafish predominately occurs during atrial systole, reflected by a low E/A ratio^48^. Given the increased reliance on atrial function in zebrafish, atrial arrhythmias are hypothesized to have a profound hemodynamic impact^48^. The E/A ratio is significantly elevated in 6 mpf female *irf8* mutants (+67.1%; p = 0.0066) (Table S3), but slightly reduced at 12 mpf relative to wild type fish (male: - 13.3%, p = 0.4928; female: -16.8%, p = 0.5474). As such, both ventricular filling time (diastasis time) and wave velocities (E/A ratio) are impacted in an age- and sex-specific manner in the *irf8* mutants.

### Loss of Irf8+ macrophages results in inflammation, epicardial activation, and fibrosis

Given the significant pathophysiology exhibited by *irf8* null hearts, we performed single cell RNA (scRNA) sequencing on hearts isolated from wild type and *irf8* mutants at 12 mpf to identify potential molecular targets contributing to these functional disruptions. We first interrogated the immune populations, as an altered immune landscape has significant ramifications on cardiac function. Based on Uniform Manifold Approximation and Projection (UMAP) for dimension reduction^55^, the immune-rich cluster in wild type and *irf8* mutant hearts was cluster 4 (Figure S5A). Compared to wild type, cluster 4 of *irf8* mutant hearts had enriched expression of several known genes involved in leukocyte infiltration and identity (Figure S5B). This included genes expressed by macrophages (*samsn1a, fcer1gl* (*p < 0.05)), T cells (*ccr9a*), neutrophils (*cybb, lyz*), and genes involved in leukocyte migration and activation (*cd74a, mmp13a* (***p < 0.001)*, lect2l* (**p < 0.01)*, card9* (*p < 0.05), and putatively *BX908782.2*^56^ (****p < 0.0001)). Additionally, *irf8* mutants lack total tissue expression of the tissue resident *csf1rb+* macrophages (Figure S5B). Since *irf8* mutant fish lack embryonic-derived macrophages while potentially acquiring recruited macrophage and monocyte populations, we quantified immune infiltration in *irf8* mutant hearts by assessing neutrophil number using fish with transgenic expression of GFP-labeled neutrophils, *Tg(mpx:GFP)*^57^. Corroborating the scRNA sequencing data, *irf8* mutants had a significant increase in cardiac neutrophil number throughout the ventricle and atrium (Figure S5C-E). An enzyme-linked immunosorbent assay (ELISA) of whole wild type and *irf8* mutant hearts also revealed that mutants had a significant increase in TNFα concentration (Figure S5F), again supporting the increased immunogenicity of *irf8* mutant hearts.

*Irf8* mutant hearts also had a unique cluster, cluster 1 (Figure 4A). Cluster 1 had enriched expression of both endothelial (*flt1, tal1, kdrl*) and mesenchymal and pro-fibrotic (*cdh11, pdgfra, fgf10a*) populations (Figure 4B). The latter populations are derived from the epicardium, the outermost layer of the heart, which undergoes epithelial-to-mesenchymal transition (EMT) in response to cardiac stress or injury (Figure 4C). These activated epicardial-derived cells (EPDCs) contribute to collagen producing myofibroblasts and fibroblast-like cells in the heart^58^ (Figure 4C). We therefore hypothesized that the presence of cluster 1 of the *irf8* mutants supports a cellular shift in molecular identity in response to local stress by both endothelial and epicardial populations.

**Figure 4.**
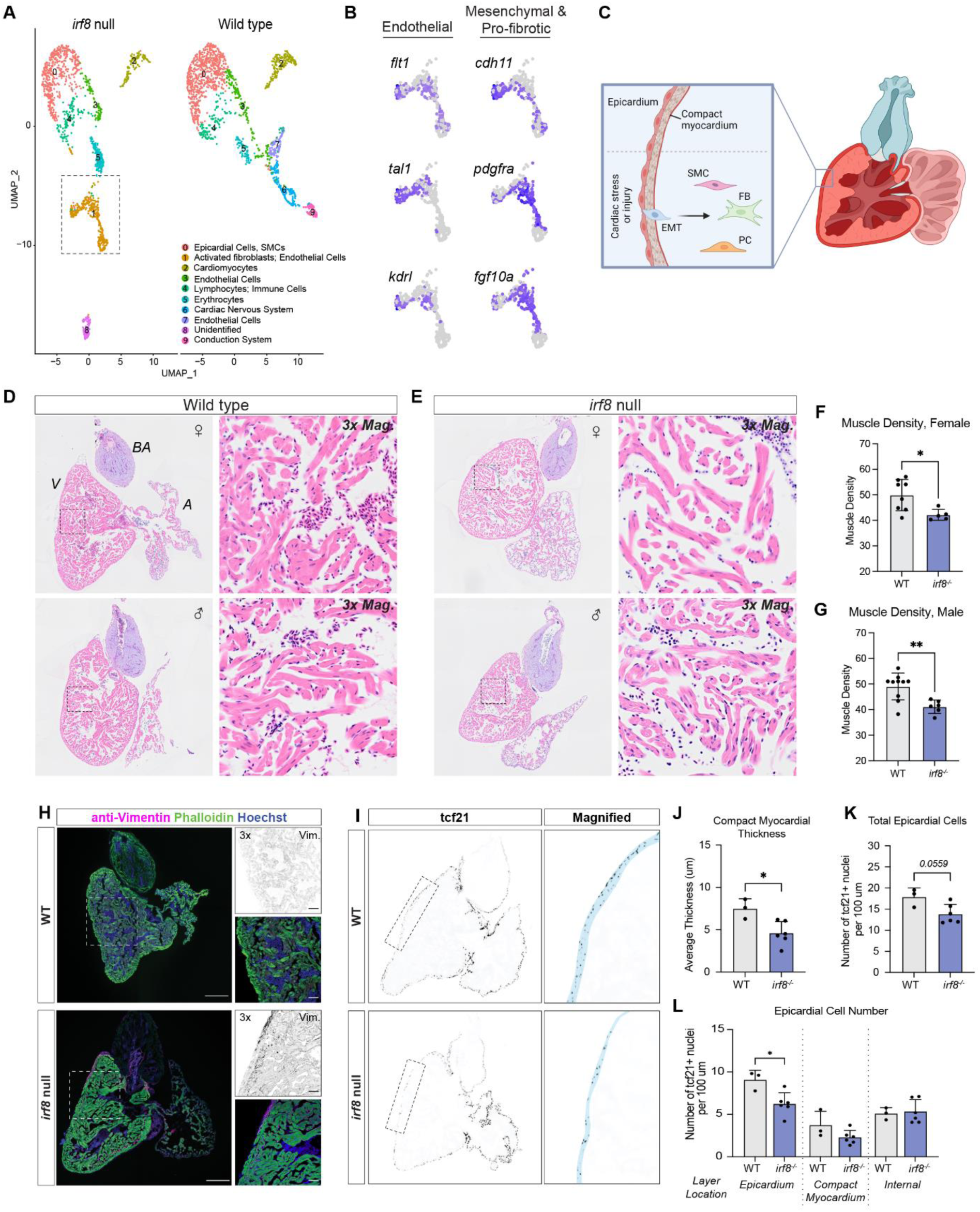
*irf8* mutant hearts are fibrotic and have epicardial abnormalities. (A) scRNA clustering by Uniform Manifold Approximation and Projection (UMAP) from wild type and *irf8* mutant hearts at 12 mpf with cluster 1 demarcated. (B) Enlarged images of cluster 1 showing expression maps of endothelial (*flt1, tal1, kdrl*) and mesenchymal and pro-fibrotic (*cdh11, pdgfrα, fgf10a*) transcripts. (C) Cartoon of the epicardium in an adult zebrafish heart. The epicardium is the outermost layer of the heart composed of squamous epithelial cells. Upon cardiac stress or injury, epicardial cells become activated and undergo epithelial-to-mesenchymal transition (EMT) to give rise to smooth muscle cells (SMCs), fibroblasts (FBs), and pericytes (PCs). Image made with Biorender. (D-E) Representative H&E sections of 12 mpf female (top) and male (bottom) wild type (D) and *irf8* null (E) hearts. (F-G) Quantification of cardiomyocyte muscle density in females (F) and males (G) at 12 mpf. Female n = 5-8 per genotype. Male n = 6-10 per genotype. Welch’s t-test. *p < 0.05. **p < 0.01. (H) Confocal micrographs of sectioned 12 mpf wild type and *irf8* mutant hearts stained for the mesenchymal marker, vimentin (Vim., magenta; gray), cardiac muscle (phalloidin, green), and nuclei (Hoechst, blue). Scale = 200 um. Images at 10x. (I) Epifluorescent images of sectioned 12 mpf wild type and *irf8* mutant hearts with transgenic expression of GFP-labeled Tcf21+ nuclei, Tg(*tcf21:nucEGFP*). The epicardial and subepicardial space (compact myocardium) is marked in blue to demonstrate thickness. (J-L) Quantification of epicardial thickness (J), total epicardial cell number (K), and epicardial cell number relative to their positioning within the epicardial and subepicardial space (L). n = 3-6 per genotype. Welch’s t-test. *p < 0.05.

Of note, endothelial populations were also present in both cluster 3 and 7 in wild type and *irf8* mutants, but *cdh11*, *pdgfra*, and *fgf10a* were either not present or minimally abundant in wild type hearts relative to *irf8* mutants (Figure S6). To address whether the abundance of mesenchymal and pro-fibrotic transcripts in *irf8* mutant hearts translated into observable differences in the tissue, we interrogated tissue architecture and mesenchymal expression of wild type and *irf8* mutant hearts directly. H&E staining of both male and female *irf8* mutant hearts revealed significantly reduced cardiomyocyte density within the ventricle (Figure 4D-G). Additionally, immunofluorescent staining for vimentin, a type III intermediate filament and cytoskeletal protein of mesenchymal cells^59^, corroborated our transcriptomic data, in that *irf8* mutant hearts had a stark increase in vimentin expression within the epicardial and subepicardial space and interstitially throughout the ventricle and atrium (Figure 4H). Abnormalities related specifically to the structure of the *irf8* mutant epicardium were also identified. The epicardium can be identified morphologically as well as by expression of epicardial markers such as Tcf21. Using a transgenic reporter line with GFP-labeled Tcf21+ nuclei to observe epicardial integrity, *Tg(tcf21:nucEGFP*)^60^, we found that Tcf21+ cells in wild type hearts formed a superficial epicardium of consistent compact myocardial thickness (Figure 4I). However, *irf8* mutants had irregular spacing between epicardial cells (Figure 4I), significantly reduced compact myocardial thickness (Figure 4J), and a near significant reduction in the number of total epicardial cells (Figure 4K). There was a significant reduction of epicardial cells on the epicardial surface of the mutant hearts, but not within the compact myocardium or internal wall adjacent to the myocardium (Figure 4L). Based on the transcriptional and immunohistological evidence of epicardial activation in the *irf8* mutants, we hypothesize that superficial epicardial cells in *irf8* mutant hearts underwent EMT, migrated inward, and adopted mesenchymal identity. Together, scRNA sequencing and immunohistological analyses support that loss of Irf8+ macrophages resulted in immune infiltration and imbalance, epicardial activation, and fibrosis.

### Irf8 mutant hearts are hyperinnervated

We next performed gene ontology analysis on marker genes in cluster 1 that were differentially expressed between clusters. Cluster 1 had enrichment of pathways predominantly related to neuron differentiation, neuron development, and axonogenesis (Figure 5A). It was shown that EPDCs may promote neurite outgrowth *in vitro*^61^, though it is not known whether EPDCs promote neurogenesis *in vivo,* especially in response to cardiac stress. We therefore assessed the single cell expression of significantly upregulated neuronal genes in cluster 1, namely *lrt2ma*, *trim46b*, and *adgrb3*, within *fgf10a*+ activated EPDCs. There was substantial co-expression of neuronal genes within activated EPDCs (Figure 5B), suggesting that epicardial activation in *irf8* mutant hearts was promoting hyperinnervation. To evaluate neuronal patterning in wild type and *irf8* mutant hearts, we performed immunostaining for the axonal projection marker anti-acetylated α-tubulin (Figure 5C-F). Confocal micrographs were skeletonized (Figure 5C’, D’, E’, F’; see ‘Methods’) and mathematically analyzed to determine the percent of axonal coverage on the ventricle and bulbus arteriosus as well as axonal compartmentalization, i.e. ‘patch’ formation, an indicator of axonal branching patterns (Figure 5C”, D”, E”, F”). The atrium was not analyzed due to collapsing tissue walls upon dissection and consequent inability to assess axons when the chamber is not dilated. At 12 mpf, *irf8* mutant hearts had a significant increase in axonal innervation on the ventricle (Figure 5G) and bulbus (Figure 5H) compared to wild type hearts. There was also a significant increase in the percent area covered in axonal patches (Figure 5I) and in total patch number (Figure 5J). When assessing the size of the patches as a percent of ventricular area, it was also determined that the patches formed on *irf8* mutants hearts were significantly larger (Figure 5K). *irf8* mutant hearts therefore were not only hyperinnervated but had aberrant branching patterns. Therefore, we affirm that, at least in part, stress responsive and activated EPDCs are aberrantly activating the cardiac nervous system contributing to the functional arrhythmias exhibited by the *irf8* mutants.

**Figure 5.**
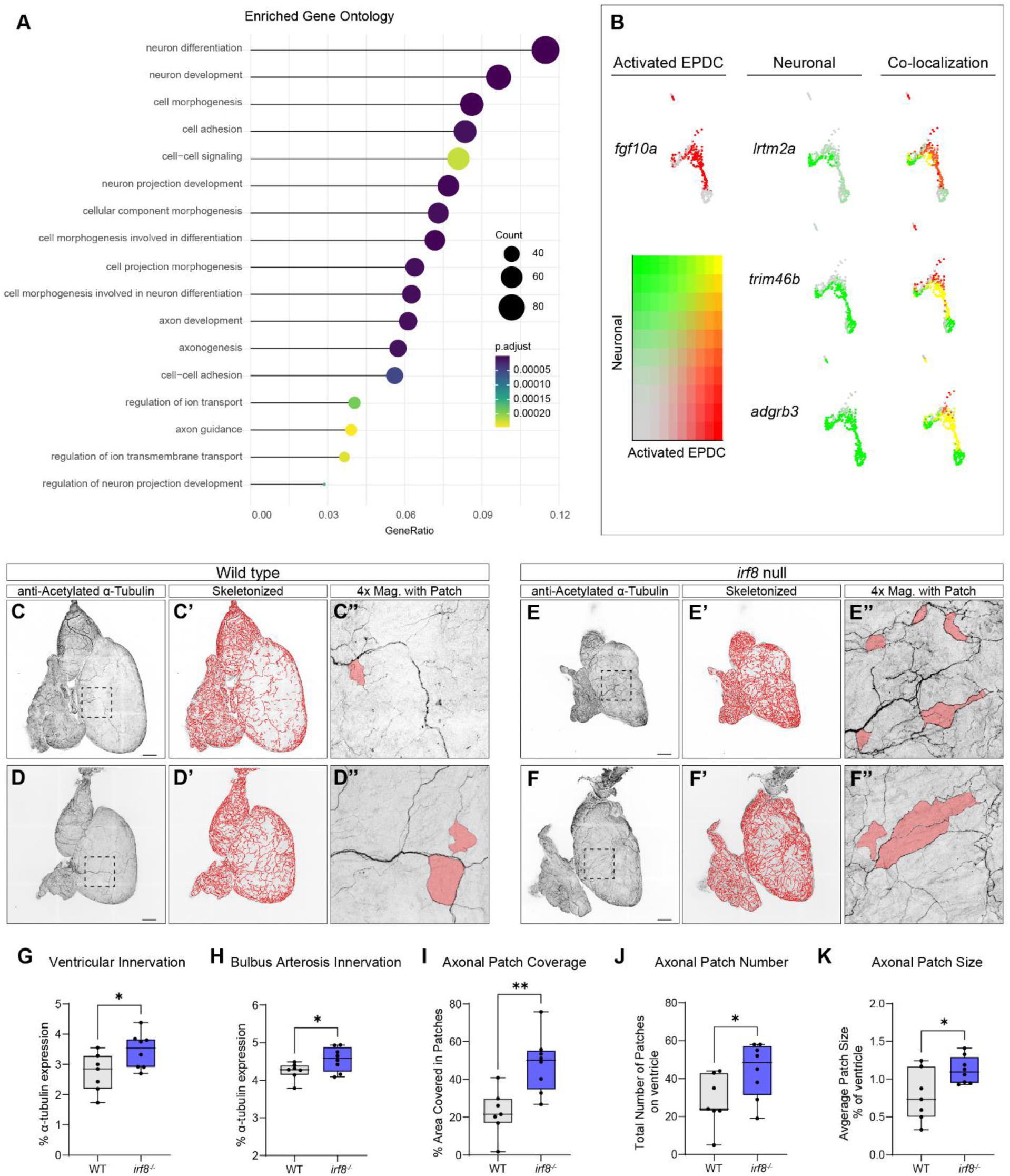
Activated EPDCs in *irf8* mutant hearts are neurogenic and hearts are hyperinnervated. (A) Enriched gene ontology analysis of cluster 1 in *irf8* mutant scRNA sequencing data at 12 mpf listing the top significantly differentially expressed genes (compared to all other clusters). (B) Co-localization analysis of activated EPDCs (*fgf10a*+) with neuronal-specific genes (*lrtm2a, trim46b, adgrb3*). (C-F’’) Confocal micrographs of 12 mpf wild type (C-D”) and *irf8* mutant (E-F”) hearts stained against anti-acetylated α-tubulin. Micrographs were skeletonized (C’, D’, E’, F’) and analyzed for axonal coverage and compartmental (patches >0.1% of ventricular area; red areas in 4x magnifications in C”, D”, E”, F”). (G-K) Quantification of 12 mpf anti-acetylated α-tubulin stained wild and *irf8* mutant hearts for ventricular innervation (G), bulbus arteriosus innervation (H), axonal patch coverage over the ventricular surface (I), number of axonal patches >0.1% of the ventricular area (J), and average axonal patch size as a percent of ventricular area (K). Scale = 200 um. Images at 10x. n = 7-8 per genotype. Welch’s t-test. *p < 0.05; **p < 0.01.

## DISCUSSION

Developments in cardiac macrophage biology have broadened our understanding of the critical cell types involved in regulating cardiac development and function. This study investigated the importance of primitive and transient definitive-derived macrophages in adult heart physiology and tissue homeostasis. Here, we report that cation influx in embryonic macrophages increased larval heart rate, highlighting the ability of macrophages to electrically modulate the developing heart. We further determined that loss of Irf8+ embryonic-derived macrophages resulted in arrhythmia, diastolic dysfunction, immune infiltration, epicardial activation, fibrosis, and hyperinnervation. This study therefore provides key insights into the subset-specific roles of macrophages in heart health and function.

This study reports several independent pathological and pathophysiological impacts in the *irf8* null zebrafish heart, which begs the question of primary versus secondary drivers of the observed phenotypes. As such, our work opens several fascinating research avenues. For example, does macrophage loss impact development of the cardiac conduction system? What is the timeline of cardiac innervation in the zebrafish heart, and when are the earliest deviations from wild type first detectable? Does the absence of embryonic-derived macrophages have a direct impact on cardiomyocyte calcium handling, or are the observed functional defects secondary to fibrosis, inflammation, and hyperinnervation? It is essential to address these questions to further underscore the importance of embryonic-derived macrophages for adult heart health. It is also worth noting that the ECG and echocardiogram data were obtained on anesthetized fish due to the limitations of immobilizing alert fish for data collection. It would be interesting to perform similar physiological assessments on alert or perhaps even active models. Despite this limitation, we observed significant impacts on several measures involving electrical conduction, hemodynamics, and chamber function across ages and sexes in anesthetized fish.

Our scRNA sequencing revealed that *irf8* mutants had a unique cluster, cluster 1, presumed to be a heterogeneous cluster of injury-responsive and pro-fibrotic cell types. Activated EPDCs of cluster 1 were also neurogenic, which has not been previously reported transcriptionally or *in vivo.* Indeed, *irf8* mutant hearts were hyperinnervated and experienced aberrant branching patterns, an endpoint not previously assessed in the context of cardiac macrophage function. Both hyperinnervation and denervation have been described in heart failure, hypertrophy, and post-myocardial infarction^62^ and have significant implications in arrhythmogenesis and prognosis^63–65^. Abnormalities in sympathetic innervation in the heart has also been shown to occur in response to hypertension^66^, heart failure^67,68^, and myocardial infarction^63^. Based on the arrhythmogenicity of adult *irf8* mutant hearts, we propose that the aberrant and excessive innervation is contributing to the observed functional disturbances. Of note, microglia, the macrophages resident to the central nervous system (CNS), are required for neuronal pruning, plasticity, and neuronal network establishment during embryogenesis^69^. Our data suggest that tissue resident macrophages may play similar roles in pruning the neural pathways that innervate peripheral systems, as well, including the intrinsic cardiac nervous system of the heart. More direct analyses of the cell types mediating cardiac innervation, such as activated EPDCs, in development and disease states are needed to clarify the mechanisms by which macrophage loss may be contributing to hyperinnervation in our study.

The lack of compaction in the H&E staining is reminiscent of noncompaction syndrome, an established cardiomyopathy with features of ‘spongy’ trabeculation, hypertrophied and hypokinetic myocardial segments, and ventricular and supraventricular arrhythmias^70,71^. Follow-up analyses of the *irf8* null hearts are warranted to assess whether our observed phenotypes are truly noncompaction syndrome or a result of reduced cardiomyocyte number, atrophy, or sarcopenia. Any of these possibilities could be a result of or result in chronic arrhythmia. Performing similar assessments in adult zebrafish using an inducible model of macrophage loss rather than genetic could address whether the observed phenotypes are progressive or could be prompted in adulthood.

The molecular mechanisms underlying the genesis of the pathological features in *irf8* mutants remain to be determined. In addition to macrophage loss, an imbalance of macrophage number and/or polarization state can also be detrimental to the heart. For example, while macrophages modulate cardiomyocyte electrical activity and promote synchronicity, increases in circulating monocytes within the heart (i.e. during hypertension, advanced age, or myocardial infarction) can promote diastolic dysfunction^17^ and impaired cardiac repair due to prolonged inflammation^72,73^. Inflammation itself is also known to increase the risk of arrhythmia, either directly through influencing cardiomyocyte ion signaling, or indirectly through extracellular matrix deposition, fibrosis, and pathological remodeling consequent to reduced oxygen availability^74–76^. Therefore, the presence of monocytic immune populations detected via scRNA sequencing in the *irf8* mutants could be contributing to the pathology seen in the hearts. Additionally, it was previously shown that neutrophilia in mice contributed to ischemia-induced ventricular arrhythmias, which was further exacerbated following macrophage loss or depletion^77^. Indeed, *irf8* mutant zebrafish embryos were previously shown to exhibit neutrophilia^37,38^, though this phenotype had not been assessed in the heart nor in adult *irf8* mutant zebrafish until now. It is still not known whether neutrophilia in *irf8* mutants results from an altered transcriptional program or compensatory upregulation in the absence of functional macrophages. It is also worth assessing the onset of neutrophilia in the *irf8* mutant hearts to determine whether early life inflammation is contributing to adult stage cardiac pathologies^78^.

Cardiac macrophages were demonstrated to be important for coronary patterning in embryonic mice^24^, AVN electrical modulation^16^, Cx43 phosphorylation and localization within the myocardium^21^, and mitochondrial homeostasis^22^. While these studies were performed in mice and were not all specific to embryonic-derived macrophages, it is plausible that disruption of any of these homeostatic functions in the zebrafish could contribute to the phenotypes observed in this study, as well.

The zebrafish *irf8* gene is an ortholog to human *IRF8*. Human *IRF8* is also expressed by several dendritic cell subsets as well as B cells, expression by which has not yet been confirmed in the zebrafish. It is also not known whether IRF8+ macrophages in humans are strictly yolk sac-derived or whether IRF8 is expressed by tissue resident macrophage populations. Considering the importance of IRF8 for tissue macrophage expression in both mice^79^ and zebrafish^37,38^, it is worth determining whether a comparable role of IRF8 in distinguishing macrophage subsets exists in humans. Interestingly, IRF8 was identified as the major risk allele in patients with systemic lupus erythematosus (SLE) who were diagnosed with coronary heart disease^80^. Studies have identified IRF8 susceptibility loci in association with other autoimmune diseases, as well, including multiple sclerosis^81^, rheumatoid arthritis^82^, and Behçet’s disease^83^. Of note, cardiac arrhythmias are also common in people with a range of autoimmune diseases^84^. It is largely accepted that autoimmune-induced arrhythmia is driven by myocardial inflammation, the presence of autoantibodies, or oxidative stress^84^. That said, given the growing evidence of macrophage maintenance in cardiac function^16,21,77^, including by this study, it is worth exploring a potential link between IRF8 susceptibility loci, the composition of immune populations in the heart, and the prevalence of arrhythmia. In conclusion, the detailed analyses herein emphasize the significance of embryonic-derived macrophages in adult cardiac homeostasis, open many exciting avenues for further research, and expand our knowledge of critical macrophage functions governing homeostatic heart health.

## ACKNOWLEDGMENTS

The authors would like to acknowledge the CardioPulmonary Vascular Biology (CPVB) Center at the Providence VA Hospital and the CPVB COBRE Service Core for their assistance in echocardiographic imaging of the zebrafish hearts. We also thank Dr. William Talbot of Stanford University for providing the *irf8^st^*^96^*^/st^*^96^ mutant line. The mouse anti-isl1/2 developed by Jessell, T.M. and Brenner-Morton, S. and the mouse anti-vimentin developed by Alvarez-Buylla, A. were obtained from the Developmental Studies Hybridoma Bank, created by the NICHD of the NIH and maintained at The University of Iowa, Department of Biology, Iowa City, IA 52242. This work was supported by the Ruth L. Kirschstein Predoctoral Individual National Research Service Award (NRSA; F31HL156460) by the NHLBI awarded to S.E.P. This work was also supported by a NIEHS Outstanding New Environmental Scientist (ONES) award, and cardiopulmonary vascular COBRE Phase II (2PG20GM103652) awarded to J.S.P. Research reported in this publication was also supported by Research Project Grants NIH NHLBI R01HL139795 (A.R.M.), R01HL163005 (A.R.M), and IDeA NIH NIGMS P20GM103652 (A.R.M.). This work was also supported by VA VHA BLR&D IK2BX002527(A.R.M.). The views expressed in this article are those of the authors and do not necessarily reflect the position or policy of the Department of Veterans Affairs or the U.S. government.

## AUTHOR CONTRIBUTIONS

Conceptualization, S.E.P. and J.S.P.; Methodology, S.E.P., C.I.O, J.A.B, and J.S.P.; Investigation, S.E.P., C.I.O, A.G., P.B., V.D.; Writing – Original Draft, S.E.P. and J.S.P.; Writing – Review & Editing, S.E.P., C.I.O., P.B., B.C., J.A.B., A.R.M., and J.S.P.; Funding Acquisition, S.E.P. and J.S.P; Resources, C.I.O., B.C., J.A.B., A.R.M., J.S.P.; Supervision, J.S.P.

## METHODS

### Zebrafish ethics statement

Zebrafish (*Danio rerio*) maintenance and experimental procedures were approved by the Brown University Institutional Animal Care and Use Committee (IACUC; 19-12-0003) adhering to the National Institute of Health’s “Guide for the Care and Use of Laboratory Animals.”

### Zebrafish husbandry

Zebrafish colonies were maintained in an aquatic housing system (Aquaneering Inc., San Diego, CA) maintaining water temperature (28.5 ± 2°C), filtration, and purification, automatic pH and conductivity stabilization, and ultraviolet (UV) irradiation disinfection. Adult and larval zebrafish were sustained in a 14:10 hour light-dark cycle (Westerfield 2000).

Adult zebrafish were placed into 1.7 L sloped spawning tanks (Techniplast, USA) 15-18 hours prior to mating. Sexes were separated by a transparent partition. Within 2 hours of light cycle onset, the partition was removed, and zebrafish were allowed to spawn for 1 hour. Embryos were collected in fresh egg water (60 mg/L Instant Ocean Sea Salts; Aquarium Systems, Mentor, OH) and placed into 100 mm non-treated culture petri dishes (CytoOne, Cat. No. CC7672-3394) until time of experiment. Embryonic and larval zebrafish were maintained at 28.5 ± 1°C in an incubator (Powers Scientific Inc., Pipersville, PA) up to 120 hours post-fertilization (hpf).

### Zebrafish and Mice Lines

The following zebrafish lines were used in this study, either independently or in combination: macrophage, *Tg*(*mpeg1:EGFP*)^40^; neurons, *Tg(elavl3:Gal4-VP16)*^85^; light-gated mono- and divalent cation channel, channelrhodopsin, *Tg(UAS:ChR2(H134R)-mCherry)s1986t*^86^; pacemaker cardiomyocytes, *Tg(isl1-hsp70l:Kaede*)^87^; cardiomyocytes, *Tg*(*isl1:Gal4VP16;myl7:mCherry*)*fh445*^88^; neutrophils, *Tg*(*mpx*:*GFP*)^57^; *irf8* mutant (TALE-NT 2.0; st96 allele)^38^; Tcf21+ epicardial cells, *Tg*(*tcf21:nucEGFP*)^60^. The following mouse line was used in this study: FVB-*Tg(Csf1r-cre/Esr1*)1Jwp/J* (stock number 019098; The Jackson Laboratory) crossed with B6.Cg-*Gt(ROSA)26Sor^tm9(CAG-tdTomato)Hze^*/J (stock number 007909; The Jackson Laboratory).

### Histology & immunofluorescent staining

Whole mount larval zebrafish, isolated adult zebrafish hearts, and isolated mouse hearts were briefly rinsed in 1x PBS and fixed in 4% paraformaldehyde (PFA; Thermo Scientific, Cat. No. AAJ19943K2) overnight at 4°C while rocking. PFA was removed and the samples washed with 3x5 minute washes with 1x PBS. Samples were processed using the appropriate experimental steps described below.

For H&E staining, fixed zebrafish hearts were stored in 70% ethanol at 4°C until time of processing. Paraffin embedding and sectioning was performed by the Brown University Molecular Pathology Core. Hematoxylin 2 (Epredia, Cat. No. 7231) and Eosin Y (Epredia, Cat. No. 7111) staining was performed using standard protocol.

For immunofluorescent staining of larval zebrafish, fixed larvae were placed in 100% cold acetone (Fisher Chemical, Cat. No. A18P) for 5 minutes at 20°C to improve permeability, followed by 3x5 minute washes with 1x PBS. Larvae were then placed in blocking solution [1x PBS with 4% bovine serum albumin (BSA; Sigma, Cat. No. A3294) and 0.6% Triton-X 100 (MilliporeSigma, Cat. No. TX15681)] overnight at 4°C while rocking. Blocking solution was removed and larvae were stained with primary antibody overnight at 4°C. Samples were washed 3x5 minutes with PBST (1x PBS with 0.6% Triton-X 100) post-primary removal, stained with secondary antibody, washed again with PBST, and stored at 20°C in VECTASHIELD antifade mounting medium (Vector Laboratories, Cat. No. H-1000) until time of imaging. For fixed adult zebrafish hearts and mouse hearts, hearts were placed in 15% sucrose (Fisher Chemical, Cat. No. S5-500) at 4°C until they fully sank to the bottom of a 1.5 mL Eppendorf tube (6-12 hours), followed by 30% sucrose at 4°C until they fully sank. Hearts were then embedded in O.C.T. (Scigen, Cat. No. 4586) and stored at -80°C until ready for cryosectioning. Cryosectioning was carried out using a Leica CM1950 cryostat set to an internal temperature around -20°C and chuck temperature around -17°C. Tissue sections were 12 um thick and adhered to Colorfrost Plus^TM^ microscope slides (Fisherbrand, 25x75x1.0mm, Cat. No. 22230892). Prior to staining, slides were placed in 1x PBS to rinse off excess O.C.T. Zebrafish hearts with transgenic expression of fluorescent markers that were not antibody stained were instead counterstained with 1:5000 Hoechst 33342 (Invitrogen, Cat. No. H3570) for 2 hours at room temperature, washed with 1x PBS, and cover-slipped (Corning, No. 1 ½, 24x50mm, Cat. No. 2980-245) with VECTASHIELD. Mouse heart sections had an extra permeabilization step of 1 hour in 0.5% Triton X-100 and 20% DMSO in 1x PBS at room temperature. Mouse sections were then washed 3x5 minutes in 1x PBS. Both zebrafish and mouse hearts undergoing antibody staining were removed from 1x PBS and blocked in PBST with 4% BSA for 1 hour at room temperature. Sections were then stained with primary antibody overnight at 4°C, washed 3x5 minutes in PBST, stained with secondary antibody for 2-4 hours at room temperature, washed again, and cover-slipped with VECTASHIELD. Slides were stored in -20°C until ready for imaging.

The following primary antibodies were used in this study: mouse anti-isl1/2 (1:100; DSHB, Cat. No. 39.4D5-concentrate); rat anti-HCN4, monoclonal (1:100; Abcam, Cat. No. ab32675); mouse anti-vimentin (1:150; DSHB, Cat. No. 40E-C-concentrate); rabbit anti-Cx45, polyclonal (1:100; BiCell Scientific, Cat. No. 00407); mouse anti-acetyl-alpha tubulin (Lys40), monoclonal (1:250; Invitrogen, Cat. No. 32-2700). The following secondary antibodies and conjugated stains were used in this study: Alexa Fluor^TM^ 405 goat anti-mouse IgG (H+L) (1:200; Invitrogen, A31553); Alexa Fluor^TM^ 488 donkey anti-rat IgG (H+L) (1:200; Invitrogen, A21208); Alexa Fluor^TM^ 633 goat anti-rabbit IgG (H+L) (1:200; Invitrogen, A21071); Alexa Fluor^TM^ 568 goat anti-mouse IgG (H+L) (Life Technologies, A11031); Alexa Fluor^TM^ 488 goat anti-mouse IgG (H+L) (1:200; Invitrogen, A11001); Alexa Fluor^TM^ 488 phalloidin (1:1000; Invitrogen, A12379).

### *In situ* hybridization via RNAscope

The RNAscope Multiplex Fluorescent Detection Kit v2 (ACDBio, Cat. No. 323100) was used to perform *in situ* hybridization to probe for *mfap4.1* (Dr-mfap4.1-O1-C3) and *pax5* (Dr-pax5-C1). The assay was performed following manufacturer instructions with the following specifications or modifications. Whole zebrafish hearts or kidneys were embedded in O.C.T. (Scigen, Cat. No. 4586) immediately following dissection and stored at -80°C until ready for cryosectioning; the “fresh-frozen sample preparation” RNAscope protocol provided by ACDBio was therefore followed. Fresh frozen tissues were cut in 12 um sections. Sections were treated with Protease IV for 15 minutes at room temperature. For fluorescent detection of the probes, the Opal 520 Reagent Pack (Akoya Biosciences, Cat. No. FP1487001KT) and Opal 620 Reagent Pack (Akoya Biosciences, Cat. No. FP1495001KT) were used (1:500 dilution in TSA Buffer).

### Genotyping

*irf8^st^*^96^*^/st^*^96^ were generated by Shiau et al. via TALEN-targeting^38^. Genotyping was carried out by fin clipping. Fins were placed directly in 50 uL of DNA lysis buffer [10 mM Tris pH 8.4 (Fisher, Cat. No. BP152-500), 50 mM KCl (Sigma, Cat. No. P9333), 1.5 mM MgCl2 (Sigma, Cat. No. M0250), 0.30% Tween-20 (Fisher, Cat. No. BP337), 0.30% Igepal/NP-40 (Sigma, Cat. No. 56741)] and incubated for 20 minutes at 94°C and cooled to 55°C. Then, 5 uL of 10 mg/mL of proteinase K (Millipore Sigma, Cat. No. 70663-4) was added and samples were incubated for 1 hour at 55°C, 20 minutes at 94°C, and held at 4°C until the PCR reaction. All temperature specific reactions were carried out in the ProFlex PCR System (Applied Biosystems). The PCR reaction was performed with 2 uL of DNA and 18 uL of master mix [diH2O, 10x Taq Buffer (Thermo Scientific, Cat. No. EP0404), 2.5 mM dNTP (Thermo Scientific, Cat. No. R1121), 25 mM MgCl2 (Thermo Scientific, Cat. No. AB0359), 10 uM forward primer (IDT; 5’-ACATAAGGCGTAGAGATTGGACG-3’), 10 uM reverse primer (IDT; 5’-GGATGAGGACCGCACTATGTTTC-3’), DMSO (Millipore Sigma, Cat. No. D8418), and Taq DNA polymerase (Thermo Scientific, Cat. No. EP0404)]. The PCR reaction conditions were as follows: 1 minute at 95°C; 35 cycles of 30 seconds at 95°C, 45 seconds at 68°C, and 30 seconds at 72°C; then 7 minutes at 72°C; and hold at 4°C. As this *irf8* mutant has a frameshift mutation at an AvaI site, the PCR product was used for restriction digest with AvaI (New England BioLabs, Cat. No. R0152L) to identify the presence of the mutation. For the restriction digest reaction, 10 uL of PCR product and 10 uL of digest master mix (diH2O, rCutSmart Buffer, and AvaI) were combined and run on the thermocycler for 20 minutes at 37°C and cooled to 4°C. The digest reaction product was run in a 2% agarose gel containing RedSafe nucleic acid staining solution (Bulldog Bio Inc, Cat. No. 21141). The gel results were visualized using the Bio-Rad Gel Doc XR+.

### Optogenetic Modulation

Modulation of macrophages or neurons was achieved by cell-specific expression of the light-gated mono- and divalent cation channel, channelrhodopsin^86^. At 7 dpf, ChR2+ or ChR2-sibling zebrafish were anesthetized in 0.02% Tricaine-S (MS-222; Pentair Aquatic Eco-Systems, TRS5) and mounted ventral-side up in 2% low-melting agarose (Fisher Scientific, Cat. No. BP160-100) surrounded by egg water. Mounted fish were placed on the stage of the stereo microscope (Zeiss) in the dark for 10 minutes prior to recording to acclimate to the environment in which the recording would take place. While in brightfield, the zebrafish heart was identified and recorded using the Noldus camera (Noldus) and pylon Viewer software (Basler AG, Ahrensburg, Germany). Heart activity was recorded at three stages: (i) during a 30 second period with just brightfield, (ii) during a 30 second period with brightfield and 405 nm light, and (iii) during a 30 second period with just brightfield that occurred 10 minutes after the (i) stage. The heart rate was determined at each stage and plotted using GraphPad Prism 9 (Dotmatics, San Diego, CA). Statistical analyses were achieved using two-way ANOVA.

### Optical mapping

Voltage mapping was performed on isolated 4 mpf zebrafish hearts. Zebrafish were anesthetized in 0.04% Tricaine-S (MS-222; Pentair Aquatic Eco-Systems, TRS5) and whole hearts were carefully isolated and cleaned of non-cardiac tissue and pericardial covering as to not disrupt the sinoatrial node. Hearts were briefly rinsed in 1x PBS and placed in sterile L-15 Leibovitz medium (Millipore Sigma; L5520) in a sterile 6-well plate. All optical mapping recordings were performed within 2 hours of dissection. Hearts were stained with voltage sensitive dye, 10 µM di-4-ANEPPS (Invitrogen, D1199), for 5-10 min and washed with media solution to avoid background fluorescence. Just prior to imaging, hearts were moved to a well containing 15 µM (-)-blebbistatin (Millipore Sigma, 2033911MG) for 10 minutes or until the heart ceased beating. The hearts were then placed in a glass bottom imaging dish (MatTek, 35mm petri dish with 14mm No. 1.5 coverglass, P35G-1.5-14-C) containing 15 µM (-)-blebbistatin. The temperature was maintained at 24°C with a dual temperature controller (World Precision Instruments). Fluorescence images of the voltage dye were acquired at 1K ∼ 2K frames/second using dual CMOS cameras (Ultima-L, 100x100 pixels, Scimedia, Japan) allowing for ratiometric image comparison to reduce motion artifacts^89^. The field of view was set to 2.5×2.5 mm2 (with 50×50 μm2 pixel resolution) using a custom-built optical apparatus, which included a Navitar lens (25 mm f0.95), two dichroic beam splitter boxes (Newport Corporation), and a refocusing achromatic lens (100 mm focal length, Newport Corporation). The optical mapping image traces were recorded and analyzed using custom-built software written in Interactive Data Language (IDL, L3Harris Geospatial Solutions) as described previously^90^. The activation of the action potential (AP) was measured from the maximum dF/dt and the repolarization of the AP was measured at maximum repolarization to the baseline.

### Echocardiography

Echocardiograms on adult zebrafish were performed using the VisualSonics Vevo3100 Imaging System following the guidance of the standardized echocardiography procedure for zebrafish described by Wang et al.^48^ Zebrafish were anesthetized in 0.02% Tricaine-S and transferred to an adapted container with an adhered slit surgical sponge to stabilize the fish during echo recording. The fish was positioned ventral-side up in the sponge and surrounded by Tricaine-S. Ultrasound images were acquired using a high frequency transducer (MS700D) mounted above the fish using a micro-manipulator and positioned along the long axis (LAX), i.e., midline of the fish. Recordings were captured within 6 minutes of placement in Tricaine-S to avoid effects of anesthesia on cardiac function^48^. After the recording, fish were placed in a recovery chamber of facility water and monitored. When possible, the same cohorts were recorded at both 6 mpf and 12 mpf to achieve accurate longitudinal assessments. Two-dimensional images to capture ventricular diastolic and systolic area, chamber volume, ejection fraction, fractional shortening, stroke volume, and cardiac output were obtained using B-Mode (frequency = 50 MHz; frame rate = 50 fps; field of view = 4.73x8.00 mm). Hemodynamic activity through the atrioventricular (AV) valve and ventriculobulbar (VB) valve was captured in Color Doppler and pulse-wave Doppler mode. Peak AV inflow velocity and VB outflow velocity were identified using Color Doppler mode. Recordings were ≥ 4 seconds in length and 8-10 cardiac cycles were measured for analysis to capture beat-to-beat variation. Analysis was performed using the Vevo Lab software package (VisualSonics, v5.6.1). Each cardiac cycle (8-10 per fish) was graphically represented for each hemodynamic measurement to emphasize the significant beat-to-beat variability, especially in the *irf8^st^*^96^*^/st^*^96^ mutants, that would be otherwise lost or underappreciated if cardiac cycles were averaged together. Statistical analyses were achieved using two-way ANOVA with post-hoc Sidak test for multiple comparisons.

### Electrocardiograms

Adult zebrafish were anesthetized in 0.02% Tricaine-S and transferred to an adapted container filled with an adhered slit surgical sponge to stabilize the fish during ECG recording. Recordings took place immediately upon the fish being unresponsive to touch and all measurements were completed within a 6-minute window of time. To record electrical impulses, we used the CardioPhys^TM^ ECG system (World Precision Instruments), which included a low-noise gold ECG probe, micromanipulator, bio-amplifier and ground electrode, in combination with the PowerLab 2/26 data acquisition unit (ADInstruments, PL2602). Surgical steel needle electrodes with pin plugs (29 gauge; 12 mm needle; 2 mm pin; ADInstruments, MLA1204) were connected to the gold ECG probe and lead wires were secured to stationary rods on the micromanipulator to control insertion of the needle electrodes into the zebrafish. For recording, the positive electrode was inserted just anterior to the ventricle in the upper left quadrant of the pericardial space and the negative electrode was placed posterior to the atrium in the lower right quadrant of the pericardial space. Care was taken to not puncture the heart. The ground electrode was placed in the Tricaine-S bath in the container. The impulse was initially filtered by the bio-amplifier (low filter = 1 Hz; high filter = 100 Hz; gain = 1000), and then was transmitted to the PowerLab via coaxial cable. Electrical activity was analyzed using LabChart Pro (v8.1.19; ADInstruments). Raw data was captured using the following settings in LabChart Pro: Low pass = 100 Hz; single-ended; mains filter on. A second channel was used to generate digitally filtered traces using a band-pass filter (high cut-off frequency = 45 Hz; low cut-off frequency = 5 Hz). Filtered traces with improved signal-to-noise were used for ECG analysis. Thirty beats were analyzed (P duration, PR interval, QRS interval, heart rate) per sample. QT and QTc intervals were not reported due to the inconsistency of accurately and confidently capturing a T wave in adult zebrafish. For analysis, 30 beats were analyzed per fish to prevent over or under-sampling of different fish due to varying heart rates.

### Swim tunnel

The swim tunnel was constructed using a 5 x 2-inch clear acrylic tubing, adapted with removable custom 3D printed filter grids at the inlet and outlet (ABS Filament, Tiertime; total diameter = 1.86 in.; thickness = 1 in.; grid size = 0.05 in.; grid hole diameter = 0.15 in.). The purpose of the custom grid is to support laminar flow such that turbulence and variable flow velocities are not generated within the tunnel. Water for the tunnel came directly from 3 water valves from our zebrafish facility racks (Aquaneering Inc., San Diego, CA) into a 2-in. PVC pipe. A flow meter (Gardena 9188-U Water Flow Meter) was connected to the pipe with a compatible PVC adaptor to monitor inward flow velocity. Maximum flow velocity achieved with our system was approximately 12 cm/s. For experimentation, 12 mpf zebrafish were acclimated in the tunnel at 1 cm/s flow for 5 minutes. Then, fish were subjected to flow rates of 5 cm/s for 10 minutes, 9 cm/s for 15 minutes, and 12 cm/s for 30 minutes. Fish were immediately removed from the tunnel then anesthetized in 0.02% tricaine for immediate ECG measurements.

### Single cell dissociation of zebrafish hearts

Dissociation of zebrafish hearts was adapted from a previously published protocol^91^. Zebrafish hearts (4 mpf, n = 4 per sex per genotype; 12 mpf, n = 3-5 per sex per genotype) were dissected and pooled in HBSS (Sigma, Cat. No. H9394) with heparin (Sigma, Cat. No. H3393) in a sterile petri dish. After tearing apart the hearts with 30 G needles, hearts were placed in 1 mg/mL of collagenase II (Sigma, Cat. No. C6885) in HBSS and placed on a shaker at 32°C, 700 x rpm, for 30 minutes. Hearts were briefly agitated by pipetting every 10 minutes to encourage full disruption of the tissue. Samples were then centrifuged at 2000 x g for 5 minutes at room temperature. After removing the supernatant, 1 mL of TrypLE Express (1x; Gibco, Cat. No. 12604013) was added and samples were placed on the shaker at 32°C, 700 x rpm, for 15 minutes. The reaction was halted upon adding 750 uL of complete media [1 mL of 20 % fetal bovine serum (Gibco, Cat. No. 10437028) into 4 mL of HBSS]. Each sample was then strained through a sterile 70 um filter (pluriSelect, Cat. No. 43-10070-50) into a 15 mL conical tube and rinsed until a total of 5 mL of complete media was used per sample. Samples were centrifuged in a rotating angle centrifuge (Beckman Coulter) at 2000 x g for 5 minutes. The supernatant was discarded, and the pellet was resuspended in 1 mL of fresh complete media and spun at 2000 x g for 5 minutes again. The supernatant was again discarded, and the pellets were resuspended in 500 uL of fresh complete media and strained through a 40-um filter (pluriSelect, Cat. No. 43-10040-50) into a sterile 1.5 mL Eppendorf tube. Cells were counted using an automated cell counter (Invitrogen Countess® II FL) using a 1:1 ratio of cells and trypan blue (Gibco, Cat. No. 15-250-061).

### Single cell library preparation and sequencing

To capture the transcriptome of zebrafish single cells, we utilized Seq-Well, a massively parallel, low-input scRNA sequencing platform^92^. Briefly, 30,000 cells in single cell suspension were loaded onto a functionalized polydimethylsiloxane (PDMS) array with wells preloaded with uniquely barcoded mRNA capture beads (Chemgenes; Macosko-2011-10(V+)). After cells had settled into wells, the arrays were then sealed with a hydroxylated polycarbonate membrane with a pore size of 10 nm, facilitating buffer exchange while confining biological molecules (such as DNA and RNA) within each well. Following membrane-sealing, subsequent buffer exchange permitted cell lysis and 3’ transcript hybridization to polyT on the beads. After removal of the membrane, the beads were pooled, and transcripts were reverse transcribed. The bead-bound cDNA product then underwent Exonuclease I treatment to remove excess primer before proceeding with second-strand synthesis and PCR amplification of the cDNA product. The amplified cDNA products were purified using SPRIselect beads (Beckman Coulter, Cat. No. B23317) and were run on a Fragment analyzer (Agilent Technologies) to verify the cDNA fragment sizes were between 0.7-2 kb. The cDNA were then fragmented and amplified for sequencing using the Illumina Nextera XT DNA Library Preparation kit (Illumina, FC-131-1024) using custom primers that enabled the specific amplification of only the 3’ ends. The resulting Illumina libraries were purified, quantified, and then sequenced on a NextSeq550 system using High output 75 cycle kits (Illumina) at an average of 15,000 reads per cell. Raw read data and processed UMI count matrices from all single cell experiments are publicly available through the NIH Gene Expression Omnibus database (accession number pending).

### Single cell transcriptome analysis

Raw reads were processed using version 2.3.0 of the Drop-seq pipeline, and according to the ‘Drop-seq Alignment Cookbook’, both found at http://mccarrolllab.com/dropseq/. Demultiplexed FASTQs were aligned to the *Danio rerio* reference genome (GRCz11) using STAR aligner. Individual reads were tagged with a 12 bp barcode and 9 bp unique molecular identifier (UMI) contained in Read 1 of each sequencing fragment. Following alignment, reads were grouped by the 12 bp cell barcodes and subsequently collapsed by the 9 bp UMI to generate a gene expression count matrix. Gene expression count matrices were analyzed and visualized in R (v.4.0.3) using the Seurat package (v.4.0.4)^93^. Cells with less than 300 genes and 500 UMIs detected were excluded from further analysis. Each group in the study was internally normalized to 10,000 transcripts, log transformed and regressed on the number of UMIs per cell before dimensionality reduction and clustering. We integrated the datasets using the canonical correlation analysis (‘CCA’) in Seurat to identify anchors/shared sources of variations between the mutant and wild type fish. Next, read data were scaled and analyzed via principal component analysis (PCA). The top PCs (n = 20) were dimensionally reduced via uniform manifold approximation (UMAP) and unsupervised clustering was performed at several resolutions to analyze differential gene expression.

An adjusted p-value cutoff of 0.05 was used to determine significance. Differential gene expression (DGE) analysis was done using the MAST package. For DGE between the mutant and wild type fish, a Bonferroni correction was applied to generate adjusted p-values.

### Innervation analysis

Using confocal micrographs (10x) of wild type and *irf8* mutant hearts stained with anti-acetyl-alpha tubulin, chambers of heart (ventricle, atrium, and bulbus arteriosus) were identified and segmented by tracing the respective borders in conjunction with the MATLAB segmentation app. Neuronal projections were enhanced within each image using a Hessian filter^94–96^. With the background noise suppressed, the neuronal axons were easily segmented with a global threshold of 0.1 (images normalized between 0 to 1) and unconnected components smaller than 100px were filtered out. The binary images were then skeletonized, and small unconnected segments (smaller than 25px were filtered out) and the total length of the axons were summed in each compartment separately. The length density was then determined for each compartment by dividing by the area of each compartment respectively. The length density for the whole heart was calculated by dividing the total length across all compartments by the area of the whole heart. We also quantified properties related to compartmentalization, i.e. ‘patch’ formation, within the ventricles, defined by regions which are fully enclosed by connected neuronal projections which are at least 0.1% of the area of the ventricle. To accomplish this, the skeleton was first corrected to ensure connectivity where appropriate, which was performed by manually inspecting each skeleton with a custom-made GUI. Next, the holes within the skeleton were filled, where a hole is a set of background pixels that cannot be reached by filling in the background from the edge of the image. The skeleton was subtracted from this image, thereby separating each distinct component. Pixels within each distinct component are connected to each other, but not to other components. Finally, morphological closing was performed to remove remnants of the skeleton which had endpoints within the edges of the patch. To determine the relative size of each patch, the area was divided by that of the ventricle. We determined the cumulative size of the patches, mean relative size, the standard deviation, and the number of patches within each ventricle.

### Protein isolation and TNFα concentration

To isolate protein from individual zebrafish hearts, dissected hearts were flash frozen in liquid nitrogen and stored at -80°C until the time of the assay. Hearts were homogenized in 200 uL of RIPA buffer (Thermo Scientific, Cat. No. J60629AK) and 0.15 mm RNAse-free zirconium oxide beads (Next Advance, Cat. No. ZROB015-RNA) in a bullet blender homogenizer (Next Advance). Samples were incubated on ice for 10 minutes then centrifuged at 13,000 x g for 20 minutes at 4°C. The protein, i.e. supernatant, was transferred to a sterile Eppendorf tube. Protein concentrations were determined by a BCA assay according to manufacturer instructions (Thermo Scientific Pierce Rapid Gold BCA Protein Assay Kit, Cat. No. A53226). Zebrafish TNFα concentrations in 12 mpf wild type and *irf8* mutant hearts were determined by an enzyme-linked immunosorbent assay (ELISA) according to manufacturer instructions (MyBioSource, Cat. No. MBS704369). Absorbance was measured at 450 nm using a microplate reader. A standard curve was generated using a four-parameter logistic curve calculator, the equation from which was used to determine the concentration of TNFα (pg/µg of total protein) in each sample.

### Statistical analyses and reproducibility

Each experiment was carried out in at least three independent experimental replicates. When possible, each experimental replicate was composed of siblings, such that wild type siblings could be compared to *irf8* mutant siblings. The appropriate statistical tests used for each dataset in this study were determined based on a variety of criteria thoroughly described previously^97^. All statistical analyses were performed using R and GraphPad Prism 9 (Dotmatics, San Diego, CA).

## SUPPLEMENTAL FIGURES

**Figure S1.**
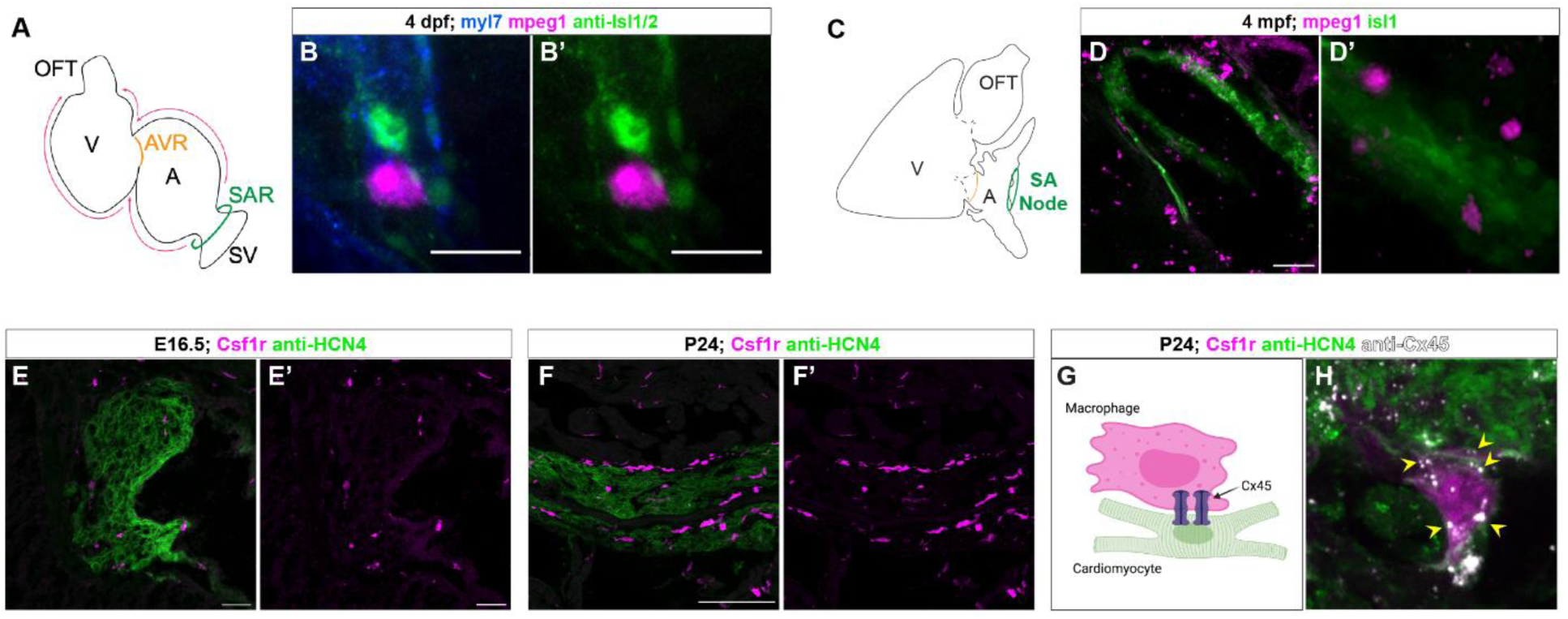
Macrophages reside at the SAN in embryonic and juvenile zebrafish and mice. (A) Schematic of a larval zebrafish heart denoting the sinus venosus (SV), sinoatrial region (SAR), atrioventricular region (AVR), atrium (A), ventricle (V), bulbus arteriosus or outflow tract (OFT), and direction of electrical propagation (pink arrows). (B-B’) Confocal micrograph of the SAR of a 4 days post-fertilization (dpf) larval zebrafish heart with transgenic expression of fluorescent cardiomyocytes (myl7+; blue), macrophages (mpeg1+; magenta), and immunostained to label pacemaker cardiomyocytes (Isl1/2+; green). Scale = 20 um. Images at 40x. (C) Schematic of the adult zebrafish heart denoting the sinoatrial (SA) node, atrium (A), ventricle (V), and outflow tract (OFT). (D-D’) Confocal micrograph of the SA node of a 4 months post-fertilization (mpf) adult zebrafish heart with transgenic expression of fluorescent macrophages (mpeg1+; magenta) and the SA node (isl1+; green). Scale = 50 um. Images at 20x. (E-E’) Confocal micrograph of the SAN (HCN4+, green) in E16.5 mice with transgenic expression of Csf1r+ macrophages (magenta). Scale = 50 um. Image at 20x. (F-F’) Confocal micrograph of the SAN (HCN4+, green) in P24 mice with transgenic expression of Csf1r+ macrophages (magenta). Scale = 100 um. Image at 20x. (G) Cartoon depiction of cardiomyocyte-macrophage coupling at the SAN via connexin45 (Cx45). (H) Confocal micrograph of the SAN (HCN4+, green) in P24 mice transgenic expression of Csf1r+ macrophages (magenta) and stained against Cx45 (white, yellow arrowheads). Image at 40x.

**Figure S2.**
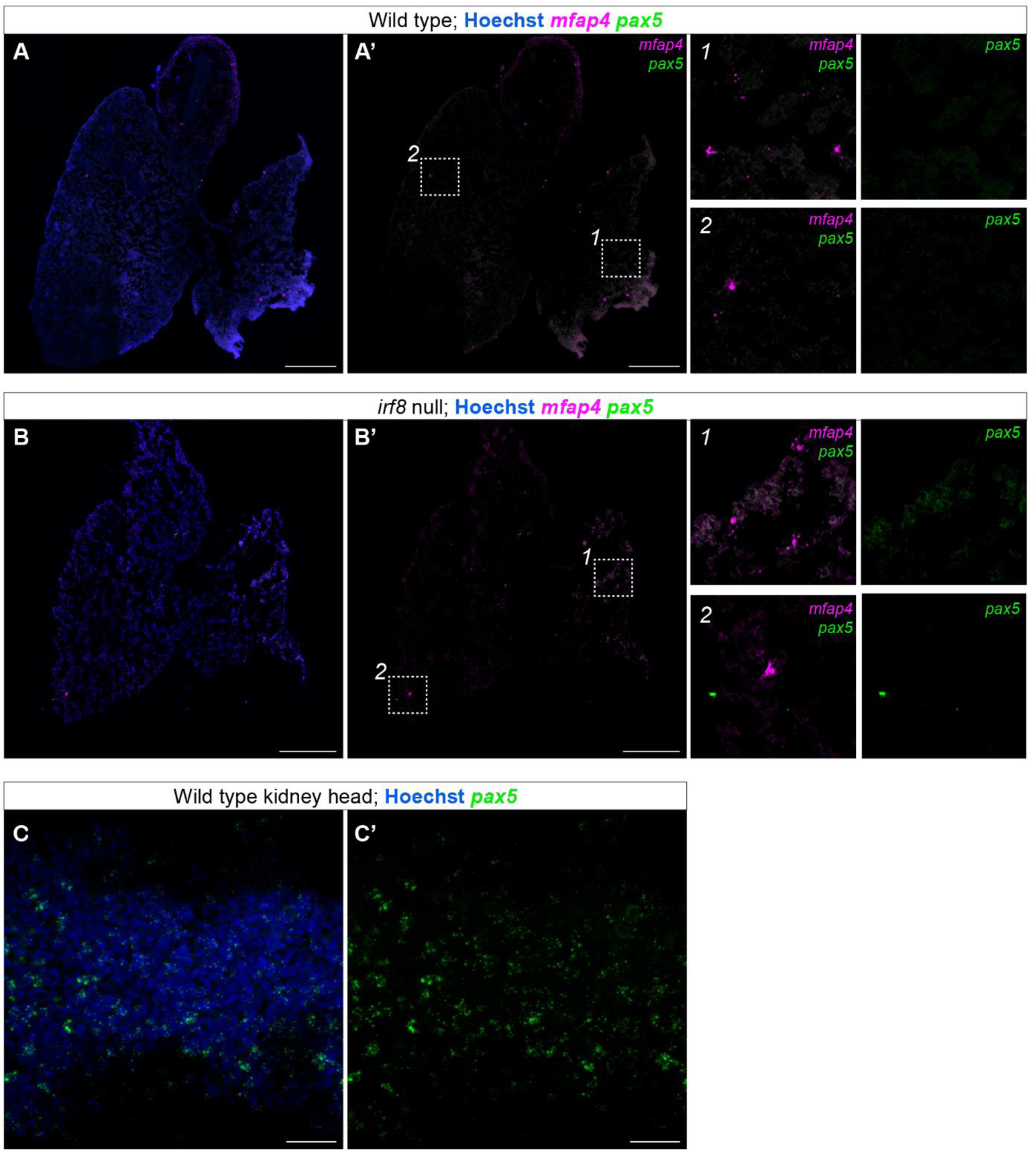
Macrophages are present in *irf8* mutant hearts. (A-B’) Confocal micrographs of 12 mpf wild type (A-A’) and *irf8* mutant (B-B’) hearts following *in situ* hybridization via RNAscope probing for macrophages (*mfap4,* magenta) and B cells (*pax5,* green). Sections were counterstained with Hoechst (blue). Boxed areas (1 & 2) in E’ and F’ are magnified by 3x. (C-C’) Confocal micrograph of an adult wild type kidney head following *in situ* hybridization via RNAscope probing for B cells (*pax5,* green). Sections were counterstained with Hoechst (blue). This was a positive control for *pax5*.

**Figure S3.**
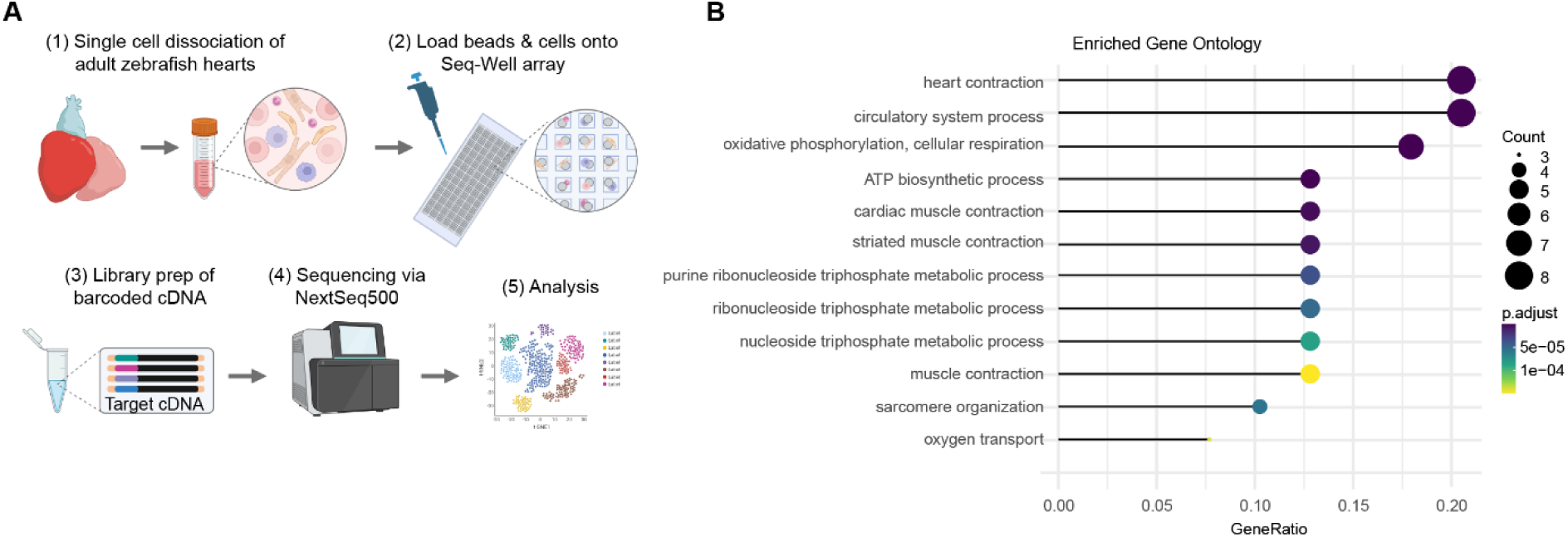
*irf8* null hearts have significant differential expression of genes associated with cardiac conduction. (A) Workflow of single cell RNA (scRNA) sequencing of adult zebrafish hearts. (B) Enriched gene ontology analysis of the significantly differentially expressed genes in 4 mpf *irf8* mutant hearts compared to wild type hearts following whole tissue analysis of scRNA sequencing data.

**Figure S4.**
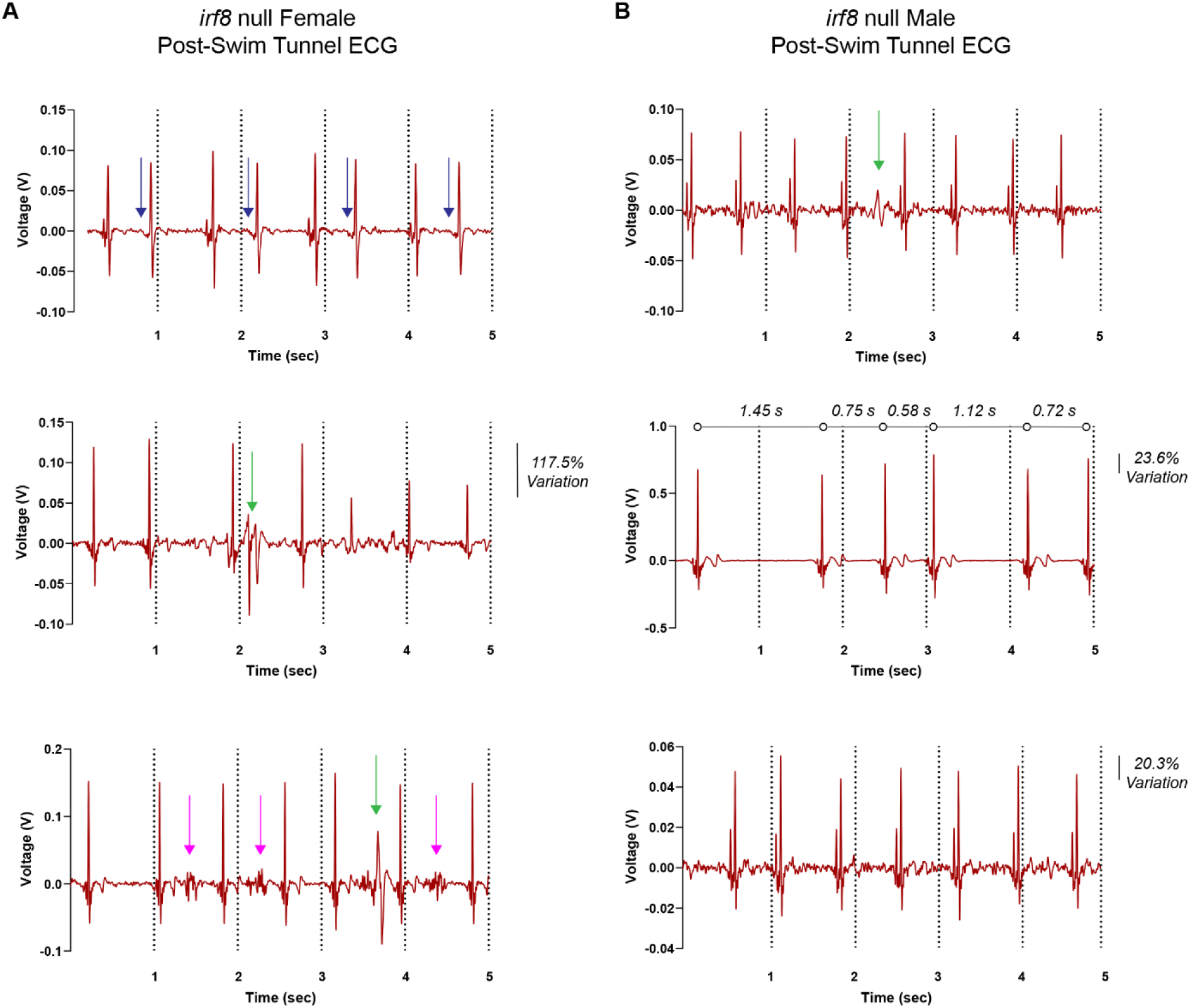
Additional examples of electrical anomalies exhibited by *irf8* null zebrafish post-swim challenge. (A) Example of electrical anomalies exhibited by 12 mpf female *irf8* null zebrafish following a 1 hr-swim challenge. Traces represent 5-second snippets of total trace. Blue arrows indicate loss or alteration of P wave consistent with premature atrial contractions in bigeminy. Green arrows represent spontaneous peaks. Pink arrows indicate flutter-like patterns. Percent variation indicates the percent change in R peak amplitude from the tallest R peak in the 5-second snippet to the lowest R peak amplitude. (B) Example of electrical anomalies exhibited by 12 mpf male *irf8* null zebrafish following a 1 hr-swim challenge. Traces represent 5-second snippets of total trace. Green arrow represents spontaneous peaks. Percent variation indicates the percent change in R peak amplitude from the tallest R peak in the 5-second snippet to the lowest R peak amplitude. Numbers between open black circles represent the time intervals between neighboring R waves.

**Figure S5.**
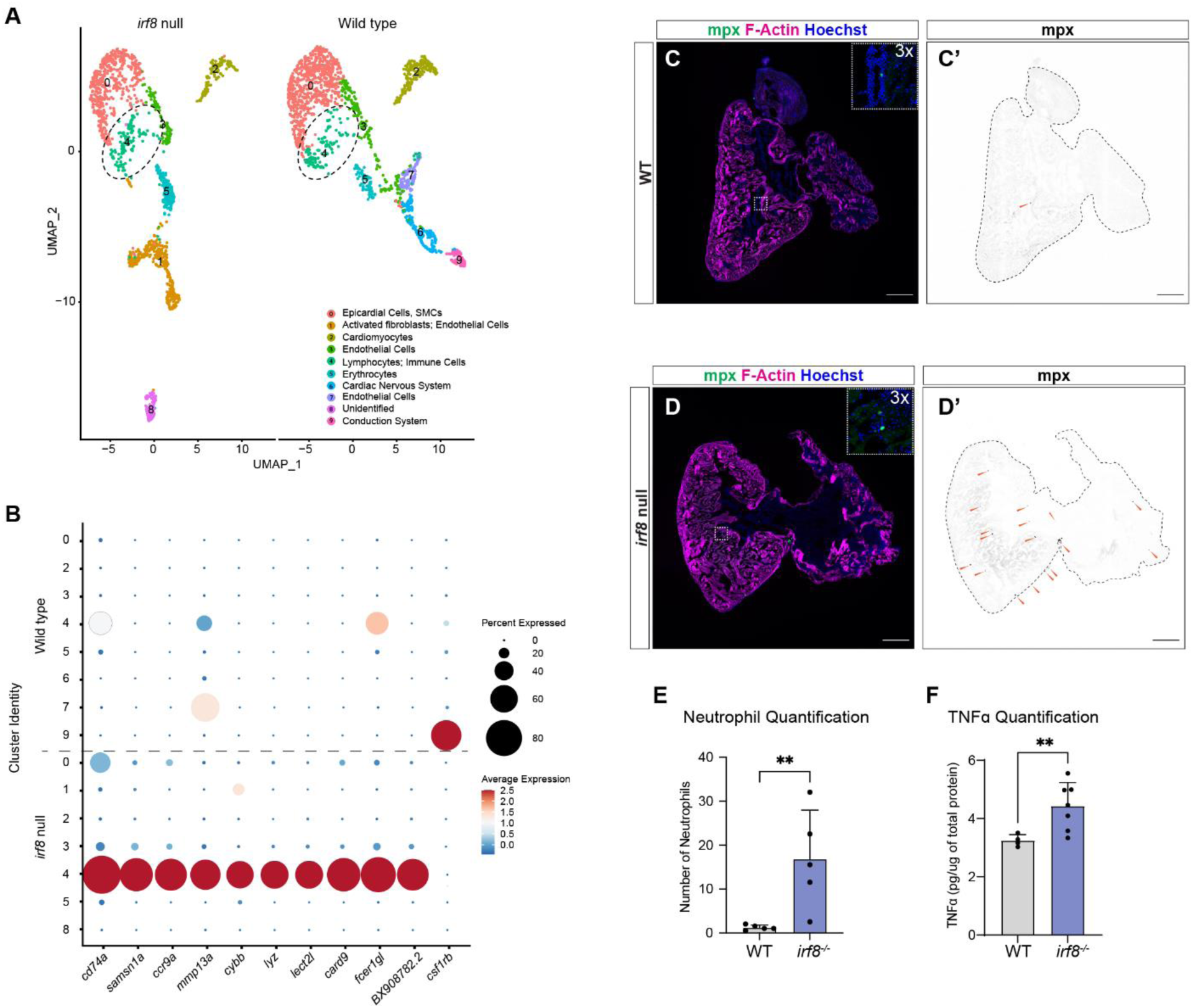
irf8 mutants have increased immune infiltration. (A) Annotated cluster map derived from scRNA sequencing data of 12 mpf wild type and *irf8* mutant hearts. (B) Dot plot displaying the expression of key genes associated with macrophages (*samsn1a, fcer1gl* (*p < 0.05)), T cells (*ccr9a*), neutrophils (*cybb, lyz*), and genes involved in leukocyte migration and activation (*cd74a, mmp13a* (***p < 0.001)*, lect2l* (**p < 0.01)*, card9* (*p < 0.05), and putatively *BX908782.2* (****p < 0.0001)). (C-C’) Confocal micrograph of a representative wild type heart with fluorescent neutrophils (green, orange arrowheads in C’) stained with phalloidin (F-Actin, magenta) and Hoechst (blue) at 6 mpf. Scale = 200 um. Image at 10x. (D-D’) Confocal micrograph of a representative *irf8* mutant heart with fluorescent neutrophils (green, orange arrowheads in D’) stained with phalloidin (F-Actin, magenta) and Hoechst (blue) at 6 mpf. Scale = 200 um. Image at 10x. (E) Quantification of neutrophils in wild type and *irf8* mutant hearts. n = 5 per genotype. Welch’s t-test. **p < 0.001. (F) ELISA for whole heart TNFα (pg/ug of total protein) in individual 12 mpf wild type and *irf8* mutant hearts. n = 5-7 per genotype. Welch’s t-test. **p < 0.001.

**Figure S6.**
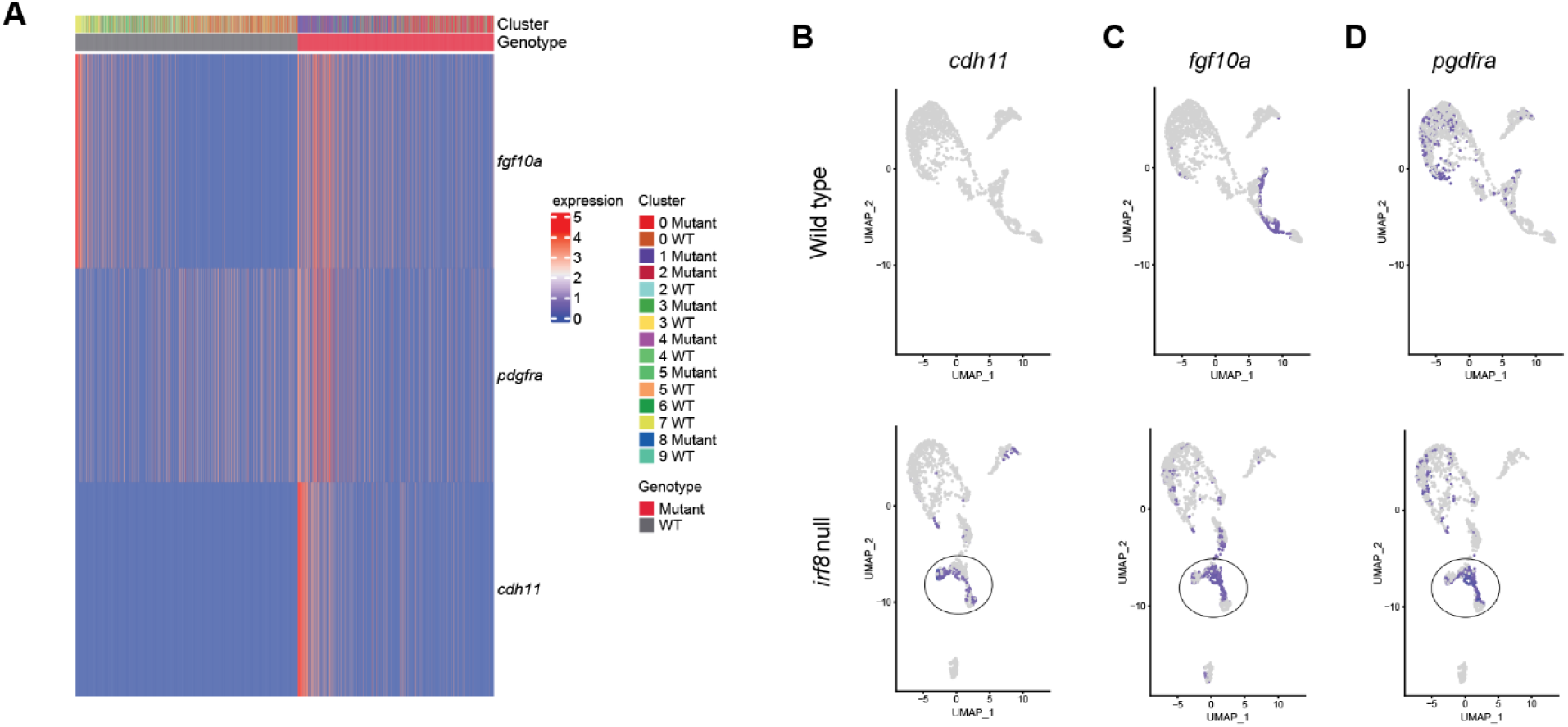
Wild type hearts have comparatively reduced expression of mesenchymal and pro-fibrotic genes relative to *irf8* mutants. (A) Heatmap of gene expression by cluster and genotype generated for the mesenchymal and pro- fibroblast genes *fgf10a, pdgfrα,* and *cdh11,* generated from scRNA sequencing data of 12 mpf wild type and *irf8* mutant hearts. (B-D) UMAP expression results of *cdh11* (B), *fgf10a* (C), and *pdgfrα* (D) in wild types and *irf8* mutants. Cluster 1 in *irf8* mutants is circled.

## Supplemental Tables

**Suppl. Table 1.** ECG results in adult unchallenged zebrafish.

|  | *irf8* WT | *irf8*<sup>st96/st96</sup> Mutant |
| --- | --- | --- |
| Heart Rate (BPM) | 68.73 ± 13.12 | **85.56 ± 15.38\*** |
| Length (cm) | 2.68 ± 0.24 | 2.62 ± 0.23 |
| Weight (g) | 0.30 ± 0.08 | 0.28 ± 0.09 |
| P Duration (msec) | 0.028 ± 0.004 | 0.027 ± 0.006 |
| PR Interval (msec) | 0.049 ± 0.010 | 0.049 ± 0.007 |
| RR Interval (msec) | 0.926 ± 0.208 | 0.738 ± 0.138 |
| QRS Interval (msec) | 0.052 ± 0.003 | 0.050 ± 0.007 |

Baseline ECG results in unexercised fish at 12 mpf. ECG measurements represent the average of 30 beats across a single recording. QT and QTc intervals are not reported here due to inconsistency of capturing and/or accurately identifying the T wave in some samples. Unpaired t-tests with Welch's correction. n = 7-9 per group, sexes combined. \*p < 0.05.

**Suppl. Table 2.** ECG results following 1-hour swim tunnel assay.

|  | Male |  | Female |  |
| --- | --- | --- | --- | --- |
|  | *irf8* WT | *irf8*<sup>st96/st96</sup> Mutant | *irf8* WT | *irf8*<sup>st96/st96</sup> Mutant |
| Heart Rate (BPM) | 67.46 ± 6.97 | 77.09 ± 16.86 | 72.48 ± 22.01 | 79.22 ± 7.87 |
| Length (cm) | 2.73 ± 0.18 | 2.74 ± 0.10 | 2.87 ± 0.12 | 2.68 ± 0.18 |
| Weight (g) | 0.31 ± 0.07 | 0.32 ± 0.05 | 0.41 ± 0.08 | 0.33 ± 0.08 |
| P Duration (msec) | 0.027 ± 0.004 | 0.027 ± 0.06 | 0.027 ± 0.006 | 0.027 ± 0.004 |
| PR Interval (msec) | 0.057 ± 0.009 | 0.053 ± 0.008 | 0.041 ± 0.015 | 0.049 ± 0.013 |
| RR Interval (msec) | 0.936 ± 0.111 | 0.863 ± 0.286 | 0.905 ± 0.291 | 0.849 ± 0.255 |
| QRS Interval (msec) | 0.052 ± 0.006 | 0.047 ± 0.008 | 0.048 ± 0.008 | 0.052 ± 0.018 |

ECG results following 1-hour swim tunnel activity at 12 mpf. No significant differences were identified between genotypes (matched sexes). ECG measurements represent the average of 30 beats across a single recording. QT and QTc intervals are not reported here due to inconsistency of capturing and/or accurately identifying the T wave in some samples. Unpaired t-tests with Welch's correction. n = 4-8 per group.

**Suppl. Table 3.**
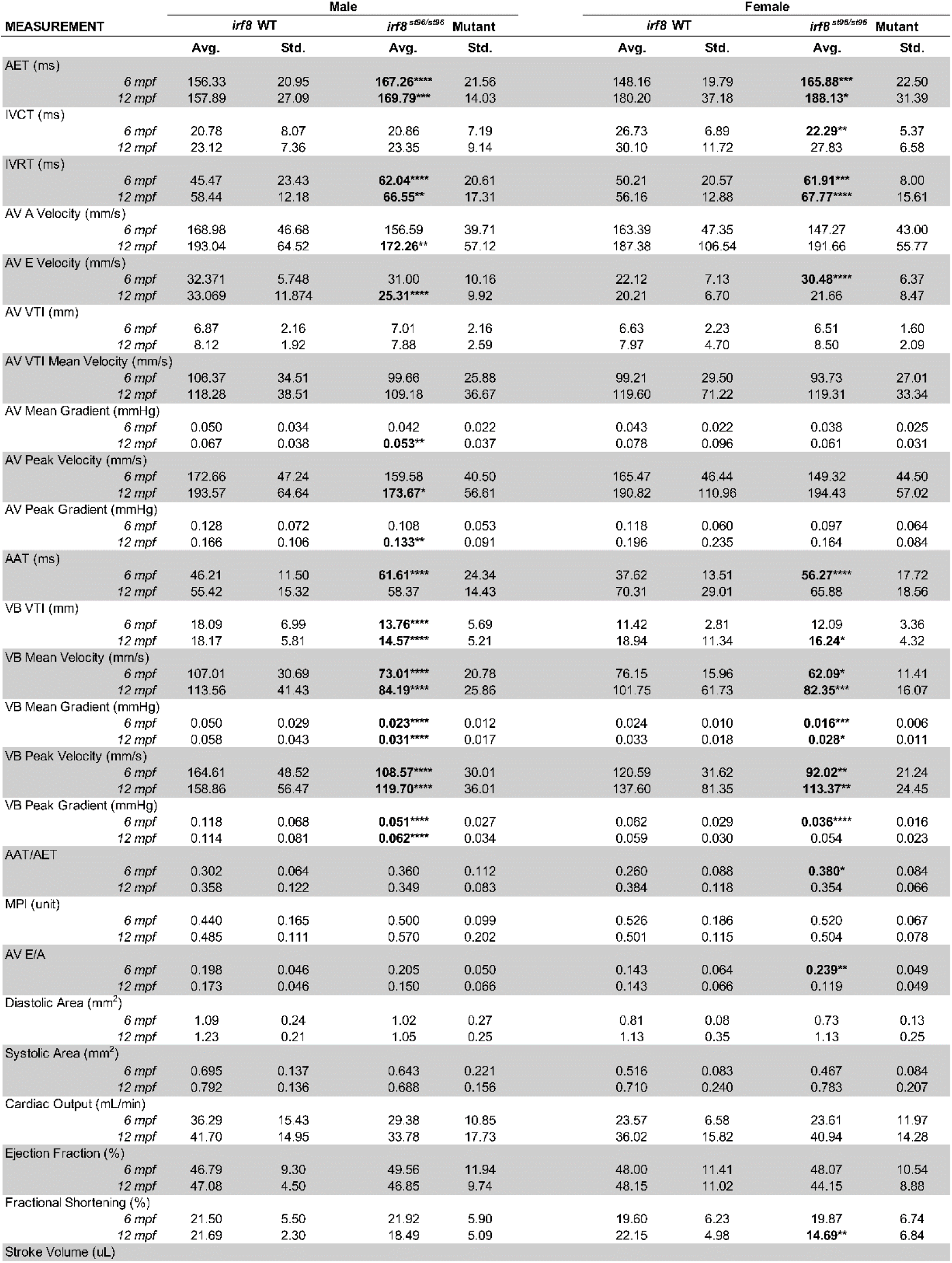
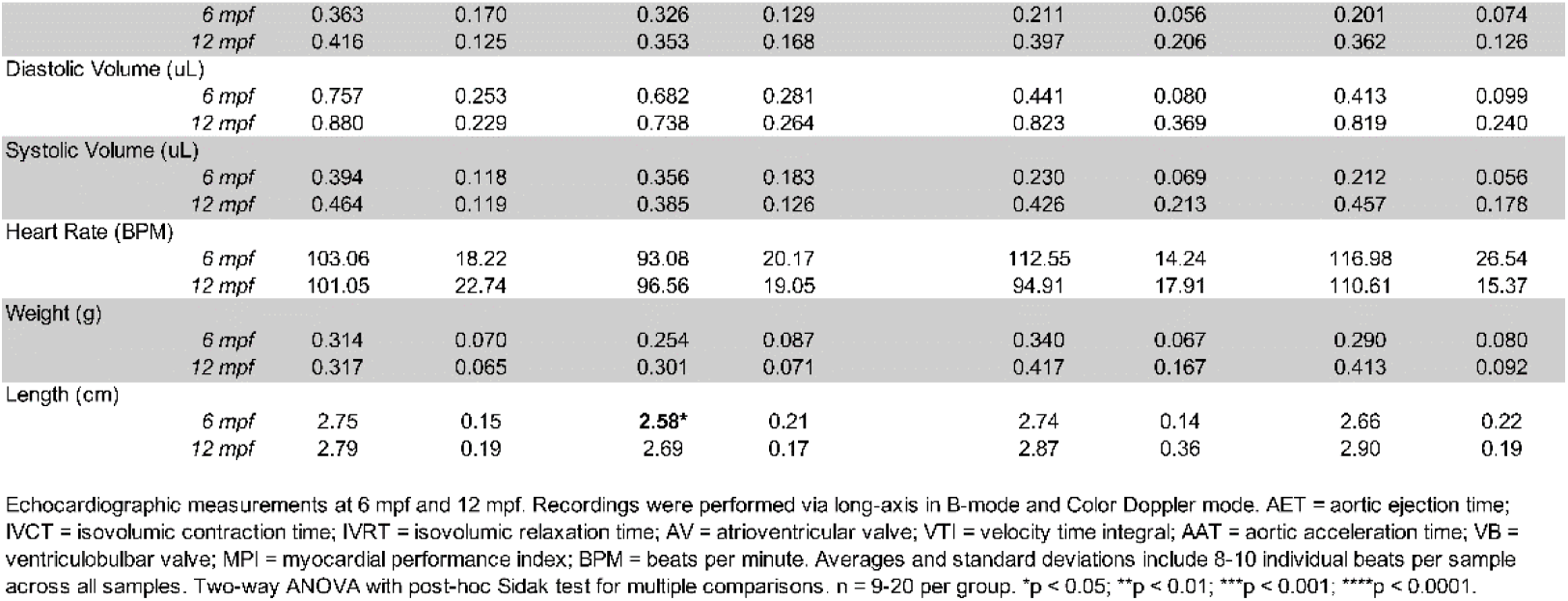
Echocardiogram results in wild type and *irf8* mutants at 6 mpf and 12 mpf.

